# A metabolic hierarchy directs cell cycle transition and morphogenesis

**DOI:** 10.64898/2025.12.20.695365

**Authors:** Shashank Agrawal, Aakash Chandramouli, Hilal Ahmad Ganie, Kavery Mohela, Infas Raheem Puthalath, Binny Mony, Urs Jenal, Siddhesh S. Kamat, Régis Hallez, Sunish Kumar Radhakrishnan

## Abstract

To optimize survival, cells must align differentiation and proliferation with metabolic status. Yet, how metabolic cues fine-tune cell cycle programs and morphogenesis remain unclear. Here, we show that cytosolic redox dynamics critically govern cell cycle transitions in *Caulobacter crescentus*. By integrating evolutionary genetics with fluxomics, we uncover temporally orchestrated shifts in core metabolic pathways that remodel cytosolic redox across the cell cycle. Early stages channel carbon flux toward unsaturated fatty-acid synthesis and a reverse-TCA to drive cytosolic oxidation-coupled G1-S transition. The later stages depend on an enhanced forward-TCA cycle that promote cytosolic reduction-driven proliferation. Strikingly, perturbing fatty-acid synthesis or reverse/forward-TCA uncouples growth from proliferation. Intriguingly, a biphasic glucose uptake program dictates the cell cycle-stage-specific metabolism. These findings reveal an unprecedented metabolic-redox circuitry coordinating morphogenesis with cell cycle progression in a carefully choreographed *ménage à trois*.

## Introduction

Metabolism provides energy and synthesizes biomolecules, which are critical to support developmental events. Beyond its fundamental role, metabolic modulations, in response to nutrient availability, is a key driving factor for development and proliferation of cells from across domains of life. For example, the decision of stem cells to proliferate or differentiate is largely dependent on the metabolic state of the cell^1^. Likewise, in the intracellular bacterial pathogen *Chlamydia trachomatis*, the differentiation between infectious elementary body versus the replicative reticulate body is influenced by the metabolic state of the pathogen and the host^2,3^. The regulatory influence of metabolism could be achieved through metabolic intermediates that impinge on the activity of biomolecules. For example, acetyl-CoA and lactate serve as substrates for enzymatic modifications of proteins or genes, thereby influencing cellular behaviour ^4,5^. Furthermore, in the dimorphic bacterium *Caulobacter crescentus*, the bifunctional activity of a key developmental regulator, KidO is influenced by its binding to NADH but not NAD^+^ ^6,7^. Another likely mechanism by which metabolism could influence protein activity could be through cytosolic redox. Notably, the existence of dynamic cytosolic redox transitions during the cell cycle has been demonstrated in prokaryotic cell types such as *Chlamydia* and *Caulobacter*, as well as eukaryotic cell types such as, mouse fibroblasts and human embryonic stem cells and epithelial cells^8–12^. The presence of redox oscillations in diverse cell types underscores its fundamental role. Nevertheless, the reason and the implications of these redox oscillations on cellular homeostasis, cell cycle progression or morphogenesis has remained unclear.

*Caulobacter crescentus* (henceforth *Caulobacter*), with its dimorphic cell fate and eukaryotic-like cell cycle presents an unique model to dissect the implications of cytosolic redox on morphogenesis, cell cycle and proliferation. During the cell cycle, *Caulobacter* cells differentiate to produce developmentally distinct daughter cells - a swarmer and a stalked cell. The stalked cell exists in an S-phase capable of replication and proliferation while the swarmer cell lies in a G1-like – replication incompetent – state. The purpose of the motile swarmer cell is to search for colonisable nutrient-rich conditions. Upon encountering favourable niche, the swarmer cell ultimately differentiates into a replication-competent staked cell. Thus, the morphogenetic transition of a swarmer to a stalked cell coincides with a G1 to S transition. Intriguingly, the cell cycle in *Caulobacter* is accompanied by a change in the cytosolic redox state.

Coincident with the G1 to S transition the cytoplasm of the swarmer cell starts to oxidise to reach maximal oxidation in the stalked cell at the mid-S-phase. From the mid-S-phase onwards the cytosolic redox gradually reduces towards the proliferative S to G2/M phase (Figure 1A, blue)^9^. This change in cytosolic redox influences activity of proteins such as the topoisomerase IV inhibitor, NstA. NstA ensures inhibition of the decatenation activity of topoisomerase IV as the chromosome replication progresses during the early S-phase. The activation of NstA is triggered by a cysteine disulphide-dependent dimerization that is facilitated in the oxidised compartment of the early S-phase cells^9^. These observations highlight the existence of a redox-based mechanism to control cell cycle and the influence of the cytosolic redox on the activity of proteins important for cell cycle. Nevertheless, the metabolic regulations of cytosolic redox oscillation and its implications on cell cycle progression remain unknown.

**Figure 1:**
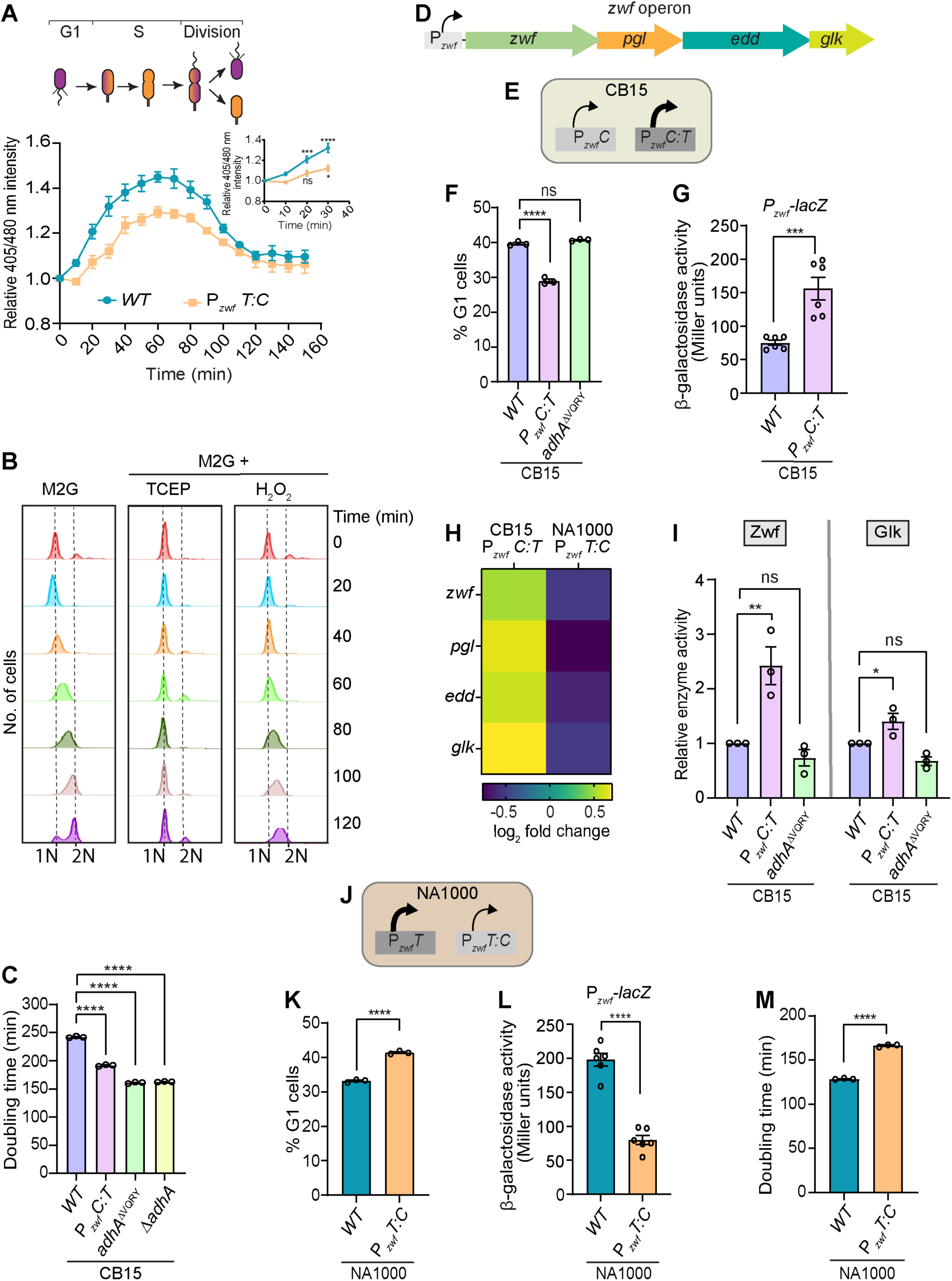
Adaptive mutation in metabolic regulators accelerate the cell cycle. **(A)** Ratio of intensity (405/480 nm) of roGFP2 during the cell cycle in a synchronized population of wild-type (*WT*) NA1000, P*_zwf_T:C* mutant. Intensity, relative to 0 min, at each timepoint is plotted (*Inset:* Ratio of intensity (405/480 nm) of roGFP2 till 30 min). Cells were grown in the presence 1mM IPTG to induce the production of *rogfp2*. **(B)** Flow cytometry profiles, representing 1N and 2N DNA content, during the cell cycle in synchronized population of *WT* NA1000 cells grown in M2G and M2G containing 20 µM H_2_O_2_ or 2.5 mM tris-(2-carboxyethyl)phosphine (TCEP). **(C)** Bar graph representing doubling time, in M2G, of *WT* CB15 and CB15 cells harbouring the mutations P*_zwf_C:T* or *adHA*^ΔVQRY^ or Δ*adHA*. **(D)** Schematic representation of *Caulobacter zwf* operon with the promoter (P*_zwf_*) along with the genes encoded in the operon. **(E)** Cartoon representing the status of *WT* P*_zwf_C* and P*_zwf_C:T* mutant promoter activity in CB15 cells. **(F)** Proportion (%) of G1 cells in *WT* CB15 and CB15 harbouring the P*_zwf_C:T* mutation or the *adHA*^ΔVQRY^ mutation. Cells were grown in M2G and analysed using flow cytometry. **(G)** β-galactosidase (LacZ) activity of the *WT* CB15 *zwf* promoter (P*_zwf_C*), or the *zwf* promoter harbouring the *C:T* mutation (P*_zwf_C:T*), fused to the *lacZ* reporter (P*_zwf_*-*lacZ*). The assay was done in exponentially growing CB15 cells. **(H)** Quantitative PCR (qPCR) analyses representing the log_2_ fold change of the transcripts encoded by the *zwf* operon in the CB15 or NA1000 cells harbouring the P*_zwf_C:T* or P*_zwf_T:C* mutations, respectively. The data represented is relative to *WT* CB15 or *WT* NA1000 transcript levels. **(I)** Enzyme activity of glucose 6-phosphate dehydrogenase (Zwf) and glucokinase (Glk) proteins, encoded in the *zwf* operon, in *WT* CB15 and CB15 harbouring P*_zwf_C:T* or *adHA*^ΔVQRY^ mutations. **(J)** Cartoon representing the status of *WT* P*_zwf_T* and P*_zwf_T:C* mutant promoter activity in NA1000 cells. **(K)** Proportion (%) of G1 cells in NA1000, and NA1000 cells harbouring the mutation P*_zwf_T:C*. Cells were grown in M2G and analysed using flow cytometry. **(L)** LacZ activity of the *WT* NA1000 *zwf* promoter (P*_zwf_T*), or the *zwf* promoter harbouring the *T:C* mutation (P*_zwf_T:C*), fused to the *lacZ* reporter (P*_zwf_*-*lacZ*). The assay was done in exponentially growing NA1000 cells.. **(M)** Bar graph representing doubling time of *WT* NA1000 and NA1000 cells harbouring P*_zwf_T:C* mutation in M2G. The data represented in C-M are from at least three independent biological replicates, ±SE. Statistical analyses were done using an ordinary one-way ANOVA with Dunnett’s multiple comparisons test in A, C, F and I, and an unpaired two-tailed t-test in G and K-M; \*\*\*\**p* < 0.0001, \*\*\**p* = 0.0002; \*\**p* = 0.0021, \**p* = 0.0332, ns = not significant. Also see Supplemental Figure S1.

Herein, we show the importance of the cytosolic redox on cell cycle progression. Moreover, using a multi-faceted approach, we have delineated the key metabolic pathways that influence cytosolic redox. We uncover an unorthodox cell cycle stage-specific metabolism, wherein when the G1 cells encounter glucose, the incoming glucose is initially favoured towards unsaturated fatty acid synthesis, triggering a peroxidation-dependent cytosolic oxidation, which in turn facilitates S-phase entry. At this stage, the TCA cycle is less abundant and an anaplerotic reaction-catalysed metabolic pathway ensures nucleotide production. As the cell cycle progresses, the forward TCA cycle is activated contributing to the conversion of oxidised cytoplasm to a reduced state. Bolstering our observations, we show that genetic perturbations of this differential metabolism imparts stage-specific arrest during cell cycle progression. Importantly, we show that this differential glucose metabolism is accompanied by an unusual biphasic uptake of glucose during the cell cycle. Together, our findings highlight an unprecedented metabolic hierarchy and its role as a frontline cell cycle checkpoint regulator.

## Results

### A genetic screen identifies metabolic regulators of cell cycle

The cytoplasmic redox of *Caulobacter* cells is dynamic during the cell cycle. The cytoplasm is reduced in the swarmer cells and becomes oxidized co-incident with the G1 to S transition to become maximally oxidized at the mid-S-phase. The reduced cytoplasm is restored towards the G2/M phase or completion of cytokinesis (Figure 1A, blue)^9^. To check whether the change in cytoplasmic redox is required for the cell cycle progression, we grew synchronized population of swarmer cells in a defined growth medium M2G having glucose as the sole carbon source, supplemented with the reducing agent Tris-(2-carboxyethyl)phosphine (TCEP) or the oxidizing agent hydrogen peroxide. Flow cytometry (FACS) analyses indicated that swarmer/G1 cells grown in the presence of TCEP were unable to enter into replicative S-phase (Figure 1B). However, in the presence of hydrogen peroxide, the G1 cells entered the replicative S-phase but could not progress in cell cycle beyond the mid-S-phase when grown in a defined growth medium such as M2G (Figure 1B). Together, these observations suggested that the cytosolic redox change during the cell cycle is crucial for the cell cycle progression, and could very well influence the growth kinetics of *Caulobacter*.

Interestingly, two isolates of *Caulobacter*, strain NA1000 and strain CB15, have different growth kinetics when glucose is used as the sole carbon source. In M2G, NA1000 has a faster growth rate when compared to CB15^13,14^. The slow growing phenotype of CB15 was attributed to a metabolic mutation^14^ and associated with a delayed G1 to S transition^13^. We wondered if the change in the cytoplasmic redox is a key contributor towards the slower cell cycle and growth in CB15. If such is the case, mutations in the components influencing the cytoplasmic redox could bestow a faster growth in CB15 cells.

To identify mutations that increase the growth rate of CB15 cells in M2G, we took advantage of an experimental evolution conducted previously^13^. Mutations accumulated in the evolved strains were identified by whole genome sequencing and the timeline of their selection was determined by sequencing the corresponding loci in intermediate timepoints. Five mutations were fixed in the evolved population #1^13^ during the first 2,000 generations (Supplemental Figure S1A). Each of these five SNPs were reconstructed individually or in combination in the ancestor CB15 strain, and their growth in M2G was measured. Strikingly, two SNPs, *CC1310^ΔVQRY^* (henceforth called as *adhA*^ΔVQRY^) and P*_zwf_C:T*, individually as well as in combination decreased the doubling time of CB15 (Figure 1C and Supplemental Table S1). Interestingly, the evolved strain selected after ∼440 generations, which harbors two SNPs (*adhA*^ΔVQRY^ and *CC0212^P77L^*) had a doubling time (157’ ± 2’) in M2G very close to the one of the reconstructed CB15 strain harboring only *adhA*^ΔVQRY^ (161’ ± 1’). On the other hand, the CB15 evolved for ∼1900 generations, which harbors three additional SNPs (P*_zwf_C:T*, *CC1158^W85*^* and *rsaA^W3G^*) had a doubling time (118’ ± 3’) lower but close to the ones of the reconstructed CB15 strains harboring either all the 5 SNPs (126’ ± 3’) or only *adhA*^ΔVQRY^ and P*_zwf_C:T* (134’ ± 1’) (Supplemental Table S1). Although CB15 *adhA*^ΔVQRY^ grew faster in M2G than CB15 P*_zwf_C:T*, the combination of both SNPs further decreased the doubling time in a CB15 background (Supplemental Table S1). In addition, these mutations had a more limited effect when minimal media containing xylose (M2X) was used instead of glucose (M2G). Complex PYE medium with or without extra glucose or xylose did not show any effect on growth of these mutants (Supplemental Table S1). Together, these results suggest that the faster growth selected in a CB15 background grown in M2G was mainly due to the additive effects of two SNPs, *adhA*^ΔVQRY^ and P*_zwf_C:T*.

### The efficiency of glucose metabolism determines the G1 lifetime

P*_zwf_C:T* is a single substitution in the promoter of the *zwf* operon (P*_zwf_*), which drives the expression of the primary glucose-metabolizing enzymes of the Entner-Doudoroff (ED) pathway including glucose-6-phosphate dehydrogenase (*zwf*), 6-phosphogluconolactonase (*pgl*), phosphogluconate dehydratase (*edd*) along with glucokinase (*glk*) (Figure 1D). The mutation converted a C at position 2268285 of the CB15 genome (AE005673.1) to a T (P*_zwf_C:T*). Growth assays indicated that CB15 cells harboring the P*_zwf_C:T* mutation were able to grow faster than the CB15 wild-type cells in M2G (Supplemental Figure S1B and Supplemental Table S1) but not in PYE or M2X (Supplemental Figure S1C and Supplemental Table S1). Furthermore, FACS based analyses showed a decreased number of G1 cells in the population of the CB15 P*_zwf_C:T* mutant when compared to CB15 wild-type (Figure 1F and Supplemental Table S2). In contrast, the CB15 *adhA*^ΔVQRY^ mutation did not alter the fraction of G1 cells (Figure 1F and Supplemental Table S2), indicating that the P*_zwf_C:T* mutation was responsible for the overall decrease in the G1 population and a faster G1 to S transition. To understand the effect of this mutation on the activity of P*_zwf_*, we used a β-galactosidase (*lacZ)*-based reporter fusion wherein P*_zwf_C* and P*_zwf_C:T* were fused to the promoter-less *lacZ* reporter gene. The P*_zwf_C:T*-*lacZ* activity was found to be significantly higher than P*_zwf_C*-*lacZ* (Figure 1G). Analyses of transcript levels by quantitative PCR (RT-qPCR), in the P*_zwf_C:T* mutant, indicated an increased abundance of all mRNAs encoded in the operon controlled by P*_zwf_* (Figure 1H). Furthermore, *in vivo* enzymatic assays indicated that the activities of Zwf and Glk were increased in cells harboring the P*_zwf_C:T* mutation (Figure 1I).

In agreement with a previous report^14^, sequence analyses revealed that the P*_zwf_* in the fast-growing NA1000 strain already harbored, at position 2294261 of its genome (NC_011916), the exact same nucleotide as the one we isolated from the evolved CB15 strain, that is a T instead of a C. Interestingly, the reversion of T to C in the P*_zwf_* of NA1000 (P*_zwf_T:C*) led to (i) a decreased P*_zwf_* activity (Figure 1L), (ii) a lowered the expression of genes encoded by the *zwf* operon (Figure 1H), (iii) a delayed the G1 to S transition (Figure 1K; Supplemental Table S2), and (iv) an increased doubling time resulting in slower growth (Figure 1M; Supplemental Figure S1D and Supplemental Table S1). Taken together, these results suggested that the efficiency of glucose metabolism could determine the G1 to S transition rate and growth in *Caulobacter* cells.

### Fatty acid abundance determines S-phase transition

The other mutation identified in our genetic screen, that significantly decreased the doubling time of CB15 in M2G, was in a gene predicted to encode an acyl-CoA dehydrogenase family protein (*adhA*). The mutation is a short deletion of 12 nucleotides (1457610-1457621; AE005673.1) leading to an in-frame deletion of the 4 amino acids VQRY at positions 72 to 75 (*adhA*^ΔVQRY^) (Supplemental Figure S1A). This mutation likely impairs the dehydrogenase activity, given that a null mutant of *adhA* (Δ*adhA*) phenocopied the *adhA*^ΔVQRY^ mutant (Figure 1C, Supplemental Figure S1B and Supplemental Table S1), indicating that a decrease of AdhA enzyme activity could increase the growth rate. Acyl-CoA dehydrogenase enzymes catalyze the first step of the β-oxidation pathway initiating the degradation of fatty acids^15,16^. Therefore, a decrease in β-oxidation pathway could lead to reduced harvest of the fatty acid pool and consequent fatty acid accumulation. Similarly, an increase in glucose metabolism through a P*_zwf_C:T* mutation, as shown above, could very well contribute towards increase in fatty acid synthesis by enhanced glucose flux through the ED pathway (Figure 2A). Indeed, our mass-spectrometry analyses of CB15 cells harboring the P*_zwf_C:T* or Δ*adhA* mutation indeed showed an increase in fatty levels when compared to the CB15 wild-type cells (Figure 2B and Supplemental Dataset 1). Moreover, the P*_zwf_C:T* mutation, with an increased level of fatty acids, was sufficient to decrease the G1 to S transition time (Figures 1F, 2B). Therefore, we wondered if the fatty acid abundance could influence the G1 to S transition. To test this, we performed mass spectrometry-based analyses to quantify the fatty acid and lipid levels, during the G1 to S transition, in NA1000 wild-type and mutant cells harboring the less active P*_zwf_T:C* promoter. Mass spectrometry-based quantifications indicated that the major fatty acid and lipid species synthesized during the G1 to S transition were significantly lower in NA1000 P*_zwf_T:C* than the native NA1000 wild-type (Figure 2C and Supplemental Dataset 3). Together, these observations indicated that glucose catabolism can indeed influence fatty acids levels and suggested that the abundance of fatty acids could influence the cell cycle progression. Next, we sought to investigate the underlying mechanism of the fatty acid-dependent control of cell cycle progression.

**Figure 2:**
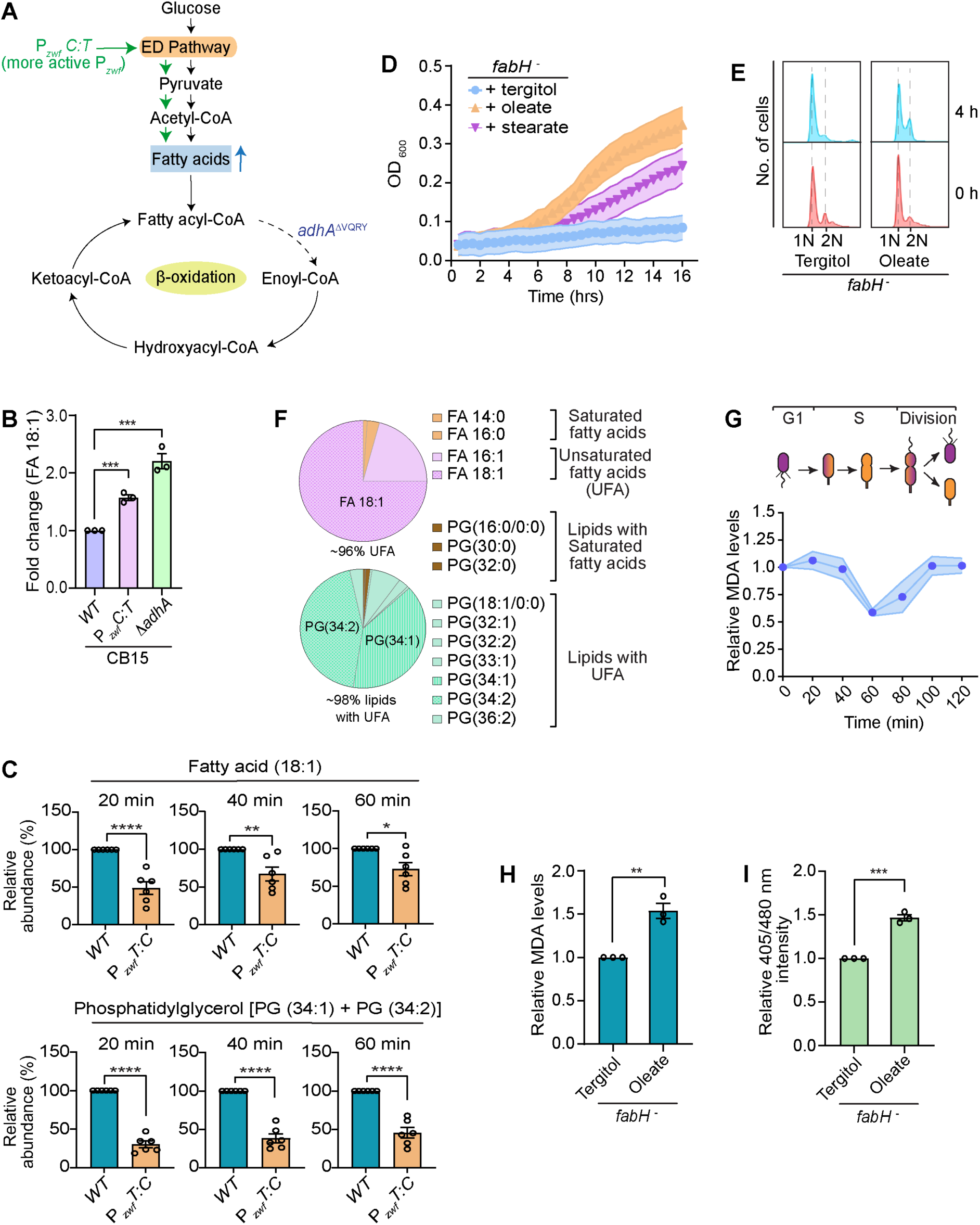
Fatty acid/lipid peroxidation triggers G1 to S transition. **(A)** Schematic representation of the glucose metabolism pathway indicating the likely effect of the P*_zwf_C:T* and *adhA*^ΔVQRY^ mutations in CB15 leading to increase fatty acid levels. Green line represents enhanced glucose metabolism in P*_zwf_C:T* while the dotted line denotes inhibition of first step of β-oxidation in *adhA*^ΔVQRY^ mutant. **(B)** Relative fold change, to wild-type (*WT*) abundance, of ^13^C-derived 18:1 species of fatty acid in CB15 cells harbouring the P*_zwf_C:T* mutation or *adhA* null mutation (Δ*adhA*). Analysis was done using exponential phase cells grown for 15 min in M2G having ^13^C_6_-glucose as the sole carbon source. Data derived from three independent biological replicates, ±SE. **(C)** Bar graph representing the % abundance, relative to *WT* levels, of ^13^C_6_-derived fatty acids (18:1) and phosphatidylglycerol (34:1 + 34:2) in NA1000 cells harbouring P*_zwf_T:C* mutation. Synchronized population of cells grown in M2G having ^13^C_6_-glucose as the sole carbon source were used to measure the abundance during early (20 min) to mid-S-phase (40 and 60 min) of the cell cycle. Data represented are from at least six independent biological replicates, ±SE, and obtained from mass spectrometry enabled ^13^C metabolic flux analyses. **(D)** Growth of *fabH* depleted cells (*fabH^-^*) in M2G medium supplemented either with 100 µM oleate, or with 100 µM stearate dissolved in 1% tergitol. Cells grown in the presence of 1% tergitol were used as control. **(E)** Flow cytometry profiles, representing 1N and 2N DNA content, in synchronized population of *fabH^-^* cells grown in M2G containing 1% tergitol or 100 µM oleate. Cells at 0h and 4h, post synchrony, were used for analyses. **(F)** Pie chart denoting the fatty acid and lipid profile in *WT* NA1000 cells. The % abundance of ^13^C_6_-derived fatty acids (FA) and phosphatidylglycerol (PG) species are represented. Data represented are from eleven independent biological replicates. **(G)** Thiobarbituric acid responsive substance (TBARS) assay denoting the relative malonaldehyde (MDA), the byproduct of fatty acid/lipid peroxidation, levels during the cell cycle in synchronized population of *WT* NA1000 cells. Data derived from three independent biological replicates, ±SE, and is relative to the initial time-point (0 min). **(H)** Relative MDA levels and **(I)** intensity ratio (405/480 nm) of roGFP2 in synchronized population of *fabH^-^* cells grown in M2G containing 1% tergitol or 100 µM oleate. *fabH^-^* cells from 60 min post synchrony were used for analyses. For D, E, H and I, depletion of *fabH* was done using the strain Δ*fabH vanA::*P*_vanA_*-*fabH*. Cells were grown in the absence of vanillate to shut down *fabH* production (see methods). Data represented in H-I are from three independent biological replicates, ±SE. Statistical analyses were done using an unpaired two-tailed t-test in B, C, H, and I; \*\*\*\**p* < 0.0001, \*\*\**p* = 0.0002; \*\**p* = 0.0021, \**p* = 0.0332, ns = not significant. Also see Supplemental Figure S2.

To gain more insights into the requirement of fatty acid synthesis during the G1 to S transition, we made use of a depletion strain for the essential 3-oxoacyl-(acyl carrier-protein) synthase III (FabH) that catalyzes the first step of fatty acid synthesis^17,18^. In the depletion strain, the only copy of *fabH* was expressed from the vanillate inducible promoter P*_vanA_* at the chromosomal *vanA* gene locus and the endogenous *fabH* gene was deleted (Δ*fabH vanA::P_vanA_-fabH)*. In this strain, the absence of vanillate leads to reduction in *fabH* expression^18^. Strikingly, cells, in which *fabH* production was depleted, were compromised in growth as well as in G1 to S transition (Figures 2D, E and Supplemental Figures S2C, D, E). Interestingly, exogenous addition of the unsaturated fatty acid, oleate (18:1), rescued growth and enabled the G1 to S transition in the *fabH* depleted cells, whereas saturated fatty acids such as stearate (18:0) could only partially rescue growth in the *fabH* depleted cells (Figure 2D). The partial growth rescue by stearate could likely be due to the desaturase activity in the cell converting stearate to oleate^19,20^. The difference in the effect of unsaturated and saturated fatty acids on growth and cell cycle progression of *fabH*-depleted cells might reflect the natural content of fatty acids in *Caulobacter* cells. In support of this, ^13^C_6_-glucose-based mass spectrometry analyses indicated that ∼96% of the total fatty acids and ∼98% of the total lipids in the *Caulobacter* cells were indeed unsaturated (Figure 2F and Supplemental Dataset 2). The most abundant fatty acid (18:1) constituted ∼75% of total fatty acids and phosphatidyl glycerol (PG) species (34:1 and 34:2) contributed to ∼85% of the total phospholipids (Supplemental Figures S2A, B and Dataset 2). Taken together, the above observations suggested that the abundance of unsaturated fatty acids/lipids is crucial for the G1 to S transition. This prompted us to investigate the mechanistic role of fatty acids in this process.

### Fatty acid/lipid peroxidation triggers cell cycle transition

It is well established that unsaturated fatty acids and lipids are prone to peroxidation^21^, which could very well influence the cellular redox. Therefore, we analyzed the fatty acid/lipid peroxidation during the cell cycle in synchronized population of NA1000 wild-type cells. Surprisingly, our results indicated that the fatty acid/lipid peroxidation happened maximally co-incident with the G1 to S transition time during the cell cycle (Figure 2G). Quantification of peroxidation levels in synchronized swarmer cells before and after addition of glucose suggested that the trigger in peroxidation is coupled to nutrient-dependent cell cycle entry (Supplemental Figure S2F). Next, to understand if the unsaturated fatty acids contributed to the redox change in the cytoplasm, we measured the lipid peroxidation and cytosolic redox state in the *fabH* depleted cells, in the presence or absence of oleate. Indeed, the *fabH* depleted cells grown in M2G supplemented with oleate had enhanced lipid peroxidation and oxidized cytosol during the G1 to S transition (Figures 2H, I).

Together, the above results emphasized the essential requirement of unsaturated fatty acids for the G1 to S transition. In wild-type cells, the cytosolic oxidation happens co-incident with the synthesis of unsaturated fatty acids that undergoes peroxidation during the G1 to S transition (Figures 1A, 2C, G). Therefore, we argued, the P*_zwf_* mutations that has decreased fatty acid synthesis (Figure 2C), and a slower G1 to S transition (Figure 1K and Supplemental Table S2), should also have an influence on cytosolic redox. To test this, we measured the redox state of the NA1000 wild-type and the NA1000 P*_zwf_T:C* mutant cells using the cytosolic-redox reporter roGFP2^9,22,23^. Strikingly, at the G1 to S transition time-point, the cytosolic redox state of the P*_zwf_T:C* mutant – having a lower P*_zwf_* activity – was significantly lower (Figure 1A, inset). Notably, the cytosolic redox state in these cells still oscillated along the cell cycle but the redox shift was delayed and the amplitude reduced as compared to wild-type cells (Figure 1A). The reduction in the P*_zwf_* activity resulted in lower abundance/activity of key glycolytic/ED pathway enzymes, resulting in a decreased flux of glucose (Figures 1H, L, 2A). Our observations suggested that the selective glucose metabolic flux influences the cytosolic redox dynamics that drives the cell cycle progression.

Altogether, these results suggested that the glucose-flux aided synthesis of unsaturated fatty acids could enhance peroxidation-dependent cytosolic oxidation that enables G1 to S transition. Inhibition of the redox change inhibits the G1 to S transition (Figure 1B), bolstering the importance of the influence of unsaturated fatty acids on cytoplasmic redox and thereby cell cycle progression.

### Differential glucose utilization during the cell cycle

Next, we wondered if the increased peroxidation during the G1 to S transition is supported by an enhanced production of unsaturated fatty acids during the early stages of the cell cycle. To test this, we measured the total fatty acid and lipid levels in NA1000 wild-type cells, during the cell cycle, using ^13^C_6_-glucose-based mass spectrometry. Indeed, our results indicated that ^13^C_6_-glucose is fluxed to synthesize fatty acid and lipids as the cell cycle is initiated (Figures 3A, B, Supplemental Figure S2G and Dataset 2). Surprisingly, we observed bi-phasic fatty acid and lipid synthesis during the cell cycle (Figures 3A, B and Supplemental Dataset 2) especially for the most abundant 18:1 fatty acid and phosphatidylglycerol (PG) (34:2, 34:1) lipid species. The first phase of fatty acid and lipid synthesis initiated at the G1 to S transition and peaked during the early S-phase. The second wave of fatty acids and lipids synthesis occurred from the late-S-phase onwards, after a pause at the mid-S-phase (Figures 3A, B and Supplemental Dataset 2).

**Figure 3:**
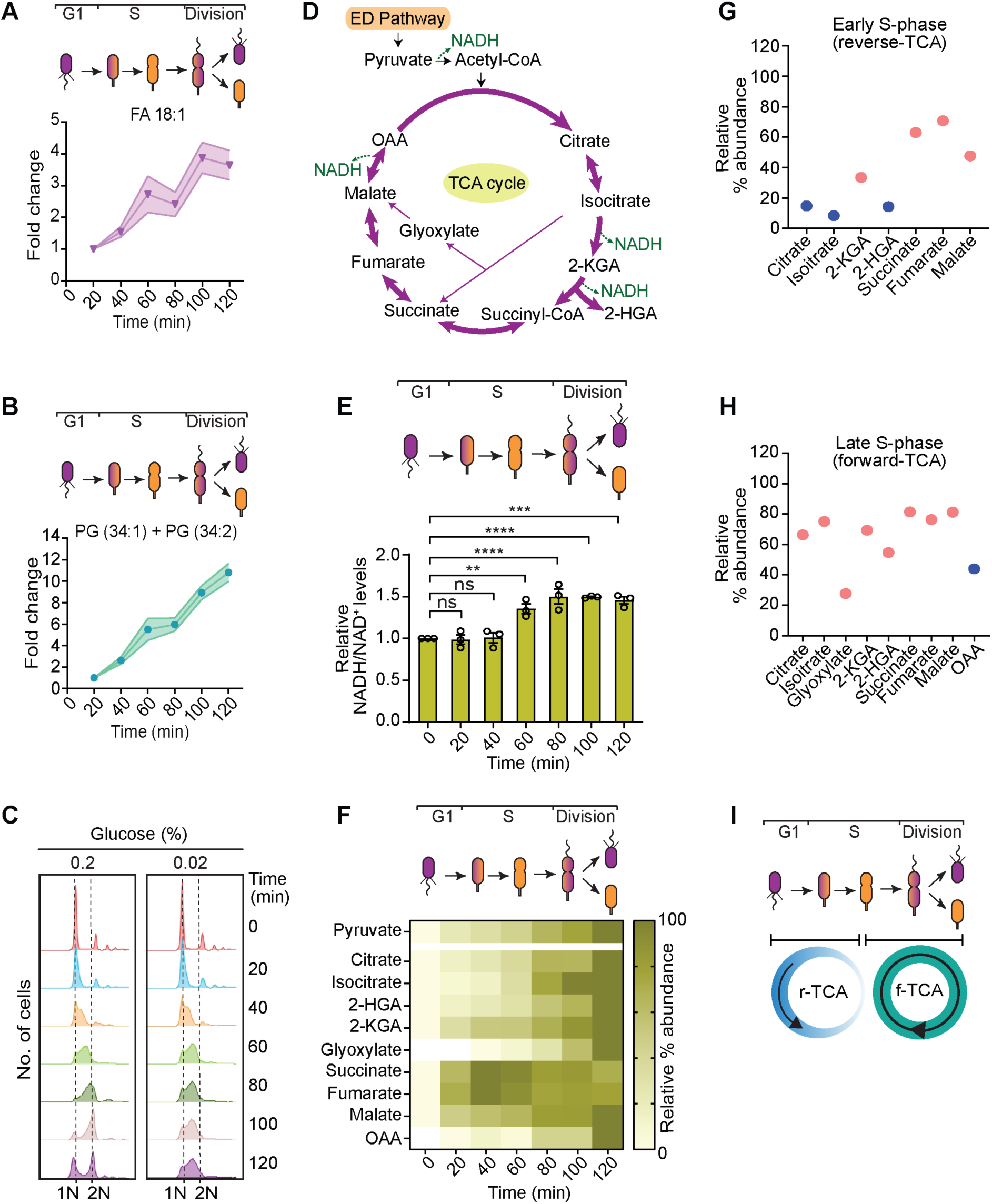
Differential glucose metabolism during the cell cycle. Mass spectrometry-based quantification of ^13^C_6_-glucose derived **(A)** fatty acid (FA 18:1) and **(B)** phosphatidyl glycerol [PG(34:1) + PG(34:2)] during the cell cycle in wild-type (*WT*) cells. Synchronized cells were grown in M2G containing ^13^C_6_-glucose as the sole carbon source and analyzed at every 20 minutes during the cell cycle. Data represented are from eleven independent biological replicates, ±SE, and is relative to the initial time-point (20 min). There were no detectable ^13^C-fatty acids/lipids at 0 min. **(C)** Flow cytometry profiles, representing 1N and 2N DNA content, during the cell cycle in synchronized population of *WT* NA1000 cells grown in M2G containing 0.2% or 0.02% glucose. **(D)** A schematic of the tricarboxylic acid (TCA) cycle and intermediates. **(E)** NADH/NAD^+^ ratio during the cell cycle in synchronized population of *WT* NA1000 cells. The experiment was performed using colorimetric assay (see methods). **(F)** Heatmap representing the abundance of TCA cycle metabolites during the cell cycle in synchronized population of *WT* NA1000 cells. Data represented are obtained from mass spectrometry enabled ^13^C_6_-glucose metabolic flux analyses. For each metabolite, the abundance (nmoles/ml/OD) was averaged from at least five independent biological replicates (Avg. abundance). To compare the abundance of a particular metabolite across the cell cycle, the Avg. abundance value was normalized to 0 to 100% (Relative % abundance). **(G, H)** Dot plots showing the comparative abundance of TCA metabolites using data derived from E. Orange dots represent metabolites having statistically significant relative abundance. **(I)** A representation of differential glucose metabolism, and the abundance of reverse and forward TCA during the cell cycle in *Caulobacter.* Statistical analyses were done using an ordinary one-way ANOVA with Dunnett’s multiple comparisons test in E, G-H; \*\*\*\**p* < 0.0001, \*\*\**p* = 0.0002; \*\**p* = 0.0021, ns = not significant. (*Abbreviations: OAA: oxaloacetic acid, 2-KGA: α-ketoglutaric acid, 2-HGA: hydroxyglutaric acid*). Also see Supplemental Figure S3.

The bi-phasic synthesis of fatty acids prompted us to wonder the reason leading to such a differential metabolism. As glucose is the only source of carbon for fatty acid synthesis in M2G, we hypothesized that the bi-phasic synthesis of fatty acids/lipids is a consequence of differential glucose utilization. To test the requirement of glucose during the cell cycle, we grew synchronized population of G1 cells in minimal media supplemented with different concentrations of glucose and analyzed cell cycle progression by measuring the DNA content using flow cytometry. Our results indicated that cells grown in 0.02% glucose were able to enter into S-phase but failed to progress beyond mid-S-phase (Figure 3C). On the contrary, the cells grown in the presence of 0.2% glucose was able to successfully complete the cell cycle (Figure 3C). Synchronized G1 cells grown in the absence of glucose failed to initiate cell cycle (Supplemental Figure S3D). To test if the cell cycle arrest at mid-S-phase in the presence of 0.02% glucose is due to glucose exhaustion, we supplemented additional 0.02% glucose at 60 mins when the cells reached mid-S-phase. This additional supplementation of 0.02% glucose was not sufficient for the progression of the cell cycle beyond the mid-S-phase (Supplemental Figure S3D). This data indicated that low levels of glucose was required for the entry into S-phase while higher amount of glucose was required from mid-S-phase onwards to complete the cell cycle.

Increased glucose requirement should be coupled with increased glucose metabolism. Therefore, as an indicator for the glucose metabolism in cells, we measured the NADH/NAD^+^ ratio using colorimetric enzymatic assay. Interestingly, the NADH/NAD^+^ ratio was increased specifically from the mid-S-phase onwards (Figure 3E and Supplemental Figures S3A, B), which coincided with the higher glucose requirement during the cell cycle. NADH is a major outcome of the tricarboxylic acid (TCA) cycle (Figure 3D)^24^. This prompted us to wonder if the differential glucose metabolism and dynamic NADH/NAD^+^ levels, observed during the cell cycle, reflect a stage-specific enhancement of TCA cycle activity at later stages of the cell cycle. To investigate this, we went ahead and measured the cell cycle prevalence of the TCA cycle using mass spectrometry-based ^13^C_6_-glucose-assisted metabolomic flux analyses. We found that the abundance of early TCA cycle metabolites such as citrate, isocitrate, and glyoxylate were indeed significantly increased from the mid-S-phase onwards (Figures 3F, H, Supplemental Figure S3C and Supplemental Dataset 4). Together, the above results suggested that (i) the absence of high amount of glucose stalls the cell cycle progression at the mid-S-phase (Figure 3C and Supplemental Figure S3D) and (ii) the TCA cycle-based glucose metabolism was enhanced from the mid-S-phase onwards (Figures 3F, H, Supplemental Figure S3C and Supplemental Dataset 4). Therefore, we wondered if the enhanced TCA cycle contributes towards the cell cycle progression beyond the S-phase. To test this, we blocked the TCA cycle in NA1000 wild-type cells by depleting the essential isocitrate dehydrogenase (IDH) with the only copy of *idh* expressed from the vanillate inducible promoter P*_vanA_* at the chromosomal *idh* locus (*idh*::*P_vanA_-idh)*. The enzyme IDH catalyzes the conversion of isocitrate to α-ketoglutarate (Figure 3D) and is a major rate-limiting step in the TCA cycle^25,26^. Remarkably, depletion of IDH led to accumulation of division defective 2N cells (Figures 4A, B), indicating that the cells were unable to progress beyond S-phase when the TCA cycle was shut down. The depletion of IDH was confirmed by the presence of increased abundance of citrate and isocitrate in the IDH depleted cells (Figures 4C, D and Supplemental Dataset 5). These results suggested that an enhanced TCA cycle is a key driver of progression into the late stages of the cell cycle to promote cytokinesis.

**Figure 4:**
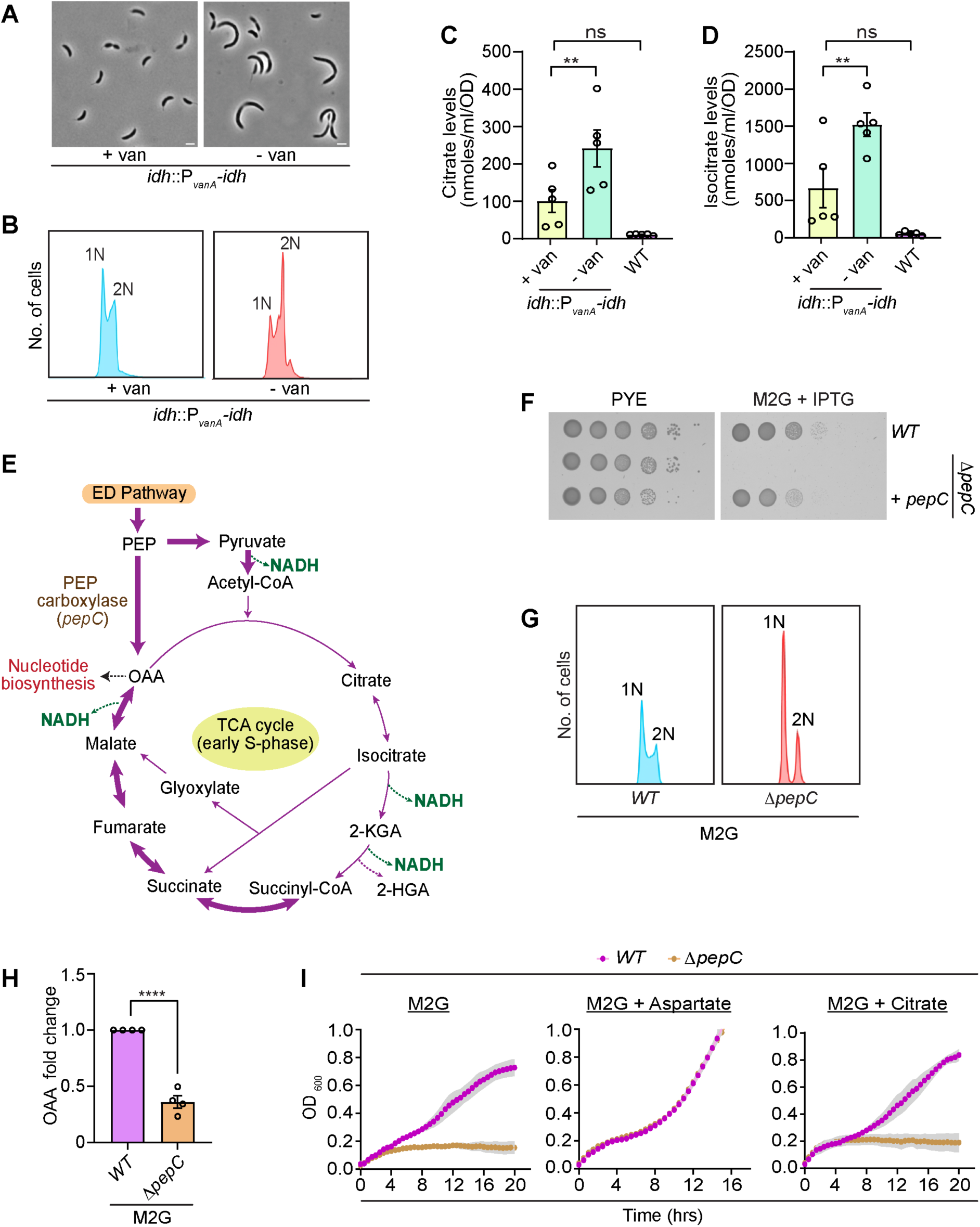
An interplay of reverse and forward TCA cycle drives cell cycle progression. **(A)** Phase-contrast micrographs and **(B)** flow cytometry profiles, representing 1N and 2N DNA content, of *idh::*P*_vanA_*-*idh* cells in the presence (+ van) or absence (- van) of vanillate. **(C, D)** Bar graphs representing the abundance of citrate and isocitrate, in wild-type (*WT*) and isocitrate dehydrogenase (*idh*) depleted cells, obtained from mass spectrometry analyses. The cells were grown in the presence of 0.5 mM vanillate to induce the production of *idh* (+ van), absence of vanillate (- van) to shut down *idh* production. Data represented are from five independent biological replicates, ±SE. **(E)** A schematic highlighting the flux of glucose through phosphoenolpyruvate (PEP) to produce oxaloacetate (OAA). This flux is mediated by the enzyme PEP carboxylase (PepC) and could enhance nucleotide biosynthesis upon S-phase entry. The part of the glucose metabolic pathway that is abundant during the early S-phase is shown using bold arrows. **(F)** Growth of *WT*, *pepC* null (Δ*pepC*) and Δ*pepC* cells ectopically expressing *pepC* from a P*_lac_* promoter on a medium copy vector (Δ*pepC* + pP*_lac_*-*pepC*). Ten-fold dilutions of the indicated strains were spotted on PYE or M2G with 1mM IPTG. **(G)** Flow cytometry profiles of *WT* and Δ*pepC* cells grown in M2G for 4h. **(H)** Mass spectrometry-based measurements of OAA levels in *WT* and Δ*pepC* cells grown in M2G for 4h. For G and H the analyses were done using cells grown overnight in a xylose containing minimal media (M2X). The cells were then washed and grown in M2G for 4h. **(I)** Graph representing the growth of *WT* and Δ*pepC* cells in M2G and M2G supplemented with either 0.2% aspartate or 0.05% citrate. Data represented in C, D, H and I are form at least three independent biological replicates, ±SE. Statistical analyses were done using an ordinary one-way ANOVA with Dunnett’s multiple comparisons test in C and D, and an unpaired two-tailed t-test in H; \*\*\*\**p* < 0.0001; \*\**p* = 0.0021, ns = not significant. Scale bar: 2 μm. (*Abbreviations: OAA: oxaloacetic acid, 2-KGA: α-ketoglutaric acid, 2-HGA: hydroxyglutaric acid*). Also see Supplemental Figure S4.

### An interplay of reverse and forward TCA cycle drives the cell cycle progression

Intriguingly, the ^13^C_6_-glucose-based metabolomic flux analyses revealed that early TCA cycle metabolites were less abundant until mid-S-phase of the cell cycle whereas, in contrast, the late TCA cycle metabolites such as succinate, fumarate and malate had an increased abundance from the early S-phase onwards (Figures 3F, G, Supplemental Figure S3C and Supplemental Dataset 4). Therefore, we wondered about the factors that could lead to the increase in the abundance of the late TCA cycle metabolites when the early metabolites are in scarce (Figures 3F, G, Supplemental Figure S3C and Supplemental Dataset 4). The accumulation of late TCA cycle intermediates might be achieved by reversible reactions since the later reactions of the TCA cycle are bidirectional (Figure 4E)^27–30^. However, for this reversible reaction to become operational, the starting metabolite of the reverse reaction – oxaloacetate – needs to be produced^31,32^. Anaplerotic enzyme-based reactions are known to replenish TCA cycle intermediates in order to meet the cellular biosynthesis demands^33^. Oxaloacetate can be synthesized from pyruvate or phosphoenolpyruvate (PEP) by the anaplerotic enzymes pyruvate carboxylase or phosphoenolpyruvate carboxylase respectively^34–36^. Analysis of the *Caulobacter* genome revealed the absence of pyruvate carboxylase. However, we found the presence of a gene (*CCNA_01560*) encoding the PEP carboxylase enzyme (PepC) in the genome. Therefore, we wondered if PepC is directly involved in the anaplerotic production of oxaloacetate. To test this, we deleted *pepC* from *Caulobacter* wild-type cells (Δ*pepC*) and found that the Δ*pepC* mutant cells were able to grow in a complex medium but not when glucose was used as the sole carbon source (Figure 4F and Supplemental S4A). This growth defect was rescued with ectopic expression of *pepC* from an inducible promoter on a plasmid (pP*_lac_*-*pepC*) (Figure 4F and Supplemental Figure S4A). Mass spectrometry analyses indicated that the Δ*pepC* cells have decreased levels of oxaloacetate (Figure 4H and Supplemental Dataset 6), suggesting that PepC indeed catalyzes the production of oxaloacetate. As the reverse TCA was abundant during the early stages of the cell cycle, it is likely that the PepC mediated anaplerotic reaction is crucial to initiate the S-phase events. If such is the case, then the Δ*pepC* cells should have increased abundance of 1N cell population in which chromosome replication has not been initiated. Indeed, FACS analyses indicated the presence of higher number of 1N cells in M2G grown Δ*pepC* mutants (Figure 4G and Supplemental Figures S4C, D).

Next, to validate the temporal importance of PepC-catalyzed reverse TCA, we made use of metabolite supplementation. It is well established that aspartate can feed into the TCA cycle through oxaloacetate. Therefore, we tested if supplementation of aspartate could rescue the growth defect of Δ*pepC* cells in M2G. Indeed, addition of 0.2% aspartate readily rescued the growth defect of the Δ*pepC* cells in M2G (Figure 4I). Furthermore, we tested if supplementation of metabolites belonging to the early part of the TCA cycle could rescue the growth defect in Δ*pepC*. Towards this, we grew Δ*pepC* cells in M2G supplemented with citrate. Interestingly, Δ*pepC* cells failed to grow in citrate supplemented M2G (Figure 4I) highlighting the cell cycle stage-specific importance of the PepC-dependent production of oxaloacetate for proper cell cycle progression.

Going forward, we wondered about the importance of reverse TCA during the early stages of the cell cycle. The PepC-catalyzed oxaloacetate production could very well operate to replenish the nucleotide pool essential for DNA synthesis upon entry into S-phase^25,37^. Therefore, we monitored nucleotide biosynthesis, which utilizes oxaloacetate-derived aspartate as precursor^38^ (Figure 5A). Using ^13^C_6_-glucose-based metabolomic flux analyses, we monitored *de novo* nucleotide biosynthesis during the early S-phase of NA1000 wild-type cells. Our analyses indicated that the indeed glucose is fluxed towards nucleotide biosynthesis during the early cell cycle stages (Figure 5B, Supplemental Figures S5A, B and Supplemental Dataset 7). Strikingly, nucleotide biosynthesis pathway-intermediates were significantly reduced in the Δ*pepC* mutant cells grown in M2G (Figure 5C and Supplemental Dataset 8).

**Figure 5:**
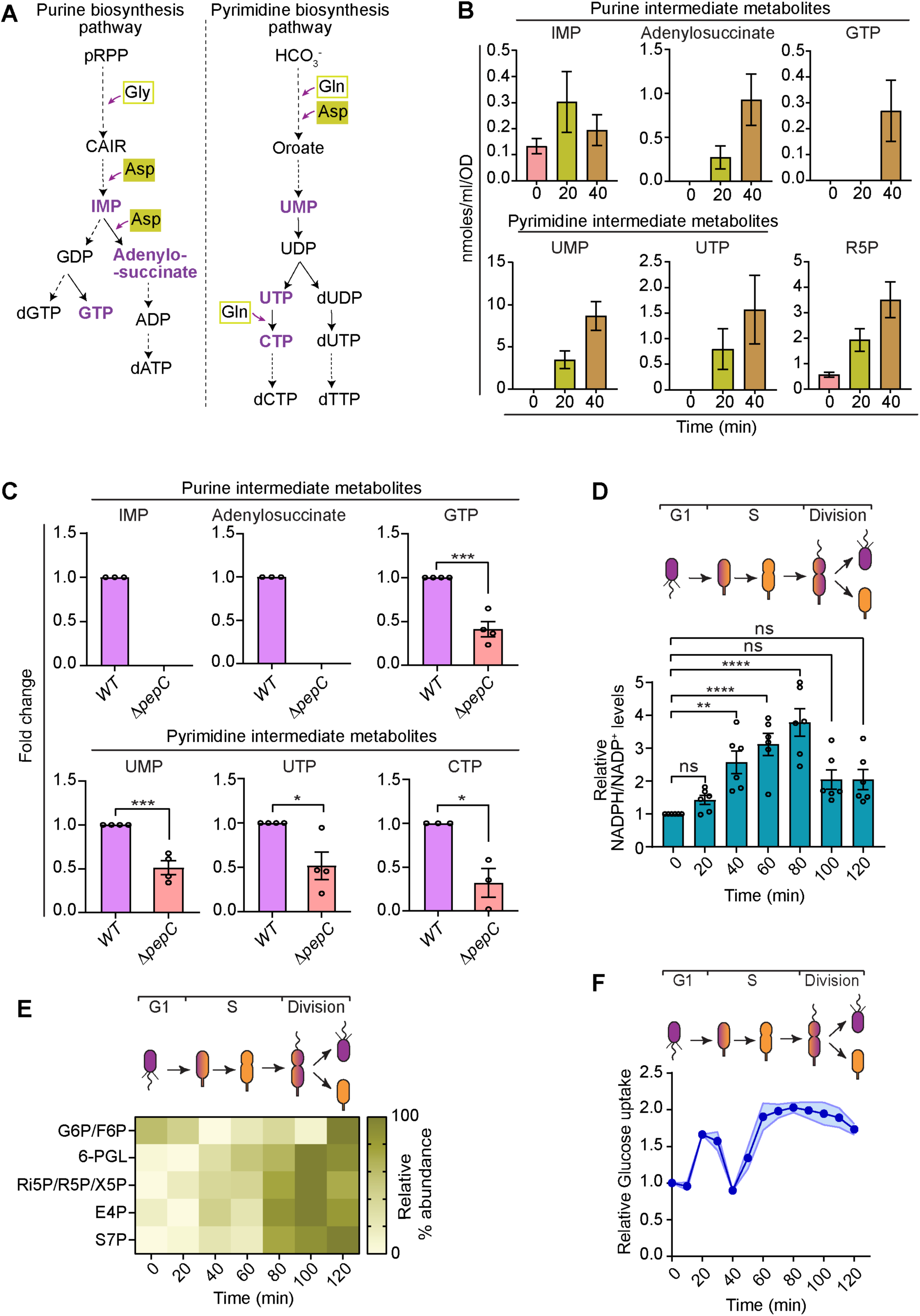
Enhanced glucose metabolism facilitates cytosolic reduction during the late S-phase. **(A)** A flow chart representing intermediates of purine and pyrimidine biosynthesis pathway. Metabolites detected in our mass spectrometry analysis is shown in purple. Empty green boxes represent amino acids glycine (Gly) and glutamine (Gln). Filled green boxes denote aspartate (Asp) derived from OAA. Dotted lines represent multi-step reactions. **(B)** Bar graph representing the abundance of ^13^C_6_-glucose derived purine, pyrimidine intermediate metabolites and ribose-5-phosphate (R5P) in synchronized population of wild-type (*WT*) NA1000 cells during early S-phase (0, 20 and 40 min). Quantification was done using mass spectrometry enabled ^13^C_6_-glucose based metabolic flux analyses. Synchronized population of swarmer (G1) cells were grown in M2G having ^13^C_6_-glucose as the sole carbon source and samples were collected at different time points as indicated. **(C)** Mass spectrometry-based measurements denoting the abundance of purine and pyrimidine intermediate metabolites in *WT* and Δ*pepC* grown in M2G for 4h. Analyses was done using cells grown overnight in a xylose containing minimal media (M2X). The cells were then washed and grown in M2G for 4h. **(D)** NADPH/NADP^+^ ratio during the cell cycle in synchronized population of *WT* NA1000 cells. The experiment was performed using colorimetric assay (see methods). Data represented are from six independent biological replicates, ±SE, and relative to the ratio at 0 min. **(E)** Heatmap representing the abundance of Entner-Doudoroff (ED) pathway metabolites during the cell cycle in synchronized population of *WT* cells. Data represented are from mass spectrometry enabled ^13^C_6_-glucose based metabolic flux analyses. For each metabolite, the abundance (nmoles/ml/OD) was averaged from at least five independent biological replicates. To compare the abundance of a particular metabolite across the cell cycle, the average abundance values were normalized to 0 to 100% (Relative % abundance). **(F)** Graph representing glucose uptake, relative to 0 min, during the cell cycle in synchronized population of *WT* cells. The analyses were done using a non-metabolizable fluorescent analog of glucose (2-NBD-glucose) and fluorescent intensity was measured at each time point as indicated. Data represented are from four independent biological replicates, ±SE. The mass spectrometry data in B, C and E are from at least five independent biological replicates, ±SE. Statistical analyses were done using an unpaired two-tailed t-test in C, and an ordinary one-way ANOVA with Dunnett’s multiple comparisons test in D; \*\*\*\**p* < 0.0001, \*\*\**p* = 0.0002; \*\**p* = 0.0021, \**p* = 0.0332, ns = not significant. (*Abbreviations: pRPP: 5-phospho-D-ribose 1-diphosphate, CAIR: N-carboxyaminoimidazole ribonucleotide, IMP: inosine monophosphate, GDP: guanosine diphosphate, dGTP: deoxy-guanosine triphosphate, GTP: guanosine triphosphate, ADP: adenosine diphosphate, dATP: deoxy-adenosine triphosphate, UMP: uridine monophosphate, UDP: uridine diphosphate, UTP: uridine triphosphate, CTP: cytidine triphosphate, dCTP: deoxy-cytidine triphosphate, dUDP: deoxy-uridine diphosphate, dUTP: deoxy-uridine triphosphate, dTTP: deoxy-thymidine triphosphate, Gly: glycine, Asp: aspartate, Gln: glutamine, G6P: glucose-6-phosphate, F6P: fructose-6-phosphate, 6-PGL: 6-phosphogluconolactone, Ri5P: ribulose-5-phosphate, R5P: ribose-5-phosphate, X5P: xylose-5-phosphate, E4P: erythrose-4-phosphate, S7P: sedoheptulose-5-phosphate)*. Also see Supplemental Figure S5.

Together, our results revealed a unique metabolic hierarchy during the cell cycle in *Caulobacter*, wherein an anaplerotic reaction triggers reverse TCA, production of oxaloacetate and synthesis of nucleotides specifically during the early part of the cell cycle. As the cell cycle progresses, the forward TCA cycle becomes enhanced, which is required for the completion of the cell cycle.

### Enhanced glucose metabolism favors cytosolic reduction

Going further, we wondered if the enhanced glucose metabolism from the mid-S-phase onwards also influences the oxidized to reduced transition of the cytosolic redox. Thanks to the activity of Zwf, glucose metabolism through ED pathway leads to the reduction of NADP^+^ to NADPH, which is known to promote a reduced environment in the cytoplasm^39^. Therefore, we went ahead and measured the NADPH/NADP^+^ ratio during the cell cycle. Interestingly, our data indicated the presence of low NADPH levels during the early stages of the cell cycle which enhanced at mid-S-phase co-incident with the onset of a reduced cytoplasm (Figure 5D and Supplemental Figure S5D). We went on to investigate if this increase in NADPH levels is coupled to increased activity of the glucose metabolizing pathways. ^13^C_6_-glucose-based metabolomic flux analyses indicated that the intermediates of the NADPH producing ED pathway are highly abundant from the mid-S-phase onwards (Figures 5E, F and Supplemental Figure S5C). Together, these results suggested that an increase in NADPH abundance at mid S-phase favors a reduced state in the cytoplasm at later stages of the cell cycle.

Next, we wondered about the reason behind increased glucose metabolism specifically from the mid-S-phase onwards. Intriguingly, the ^13^C_6_-glucose-based metabolomic flux data indicated that glucose-6-phosphate levels are decreased at around 40min into the cell cycle (Figure 5E). Glucose is phosphorylated to glucose-6-phosphate by glucokinase immediately upon uptake^40^. Therefore, a decrease in glucose-6-phosphate levels could very well indicate a decreased glucose uptake. This led us to investigate if cellular uptake of glucose is regulated during the cell cycle. To measure glucose uptake during the cell cycle, we used the non-metabolizable fluorescent analog of glucose (2-NBD-glucose). Interestingly, we observed two stages of glucose uptake during the cell cycle – the first wave of uptake happened upon the G1 to S transition and the second, more enhanced wave of uptake, occurred from the mid-S-phase onwards (Figure 5F). This result highlighted a unique phenomenon wherein the glucose uptake is regulated during the cell cycle. This observation further suggested that the dynamic glucose uptake could drive the differential glucose metabolism during the cell cycle.

## Discussion

Metabolism, a fundamental determinant of developmental processes, must be tightly integrated with cell cycle progression and proliferation to maintain cellular homeostasis. The influence of metabolism on cell cycle has been demonstrated in cell types from different domains of life. For example, the decision of a stem cell to proliferate or differentiate is largely governed by its metabolic state^1^. However, our knowledge on the implications of metabolism on cell cycle transitions, morphogenesis, and proliferation has remained poorly understood. Herein, using the dimorphic bacterial model *Caulobacter*, we unravel cell cycle stage-specific metabolic processes that influence cell cycle progression. Against the notion that core metabolism is perpetual in a cell, our results suggest the existence of dynamic metabolic processes that directly impinge on cell cycle progression and morphogenesis. We show that perturbing distinct metabolic pathways causes stage-specific arrest of cell cycle progression. Furthermore, we demonstrate that these metabolic states modulate cytosolic redox dynamics and metabolic rewiring that disrupts redox balance similarly affects cell cycle progression. Together, our findings identify a metabolism-dependent redox regulatory axis that functions as a checkpoint controlling cell cycle progression in *Caulobacter* (Figure 6).

**Figure 6:**
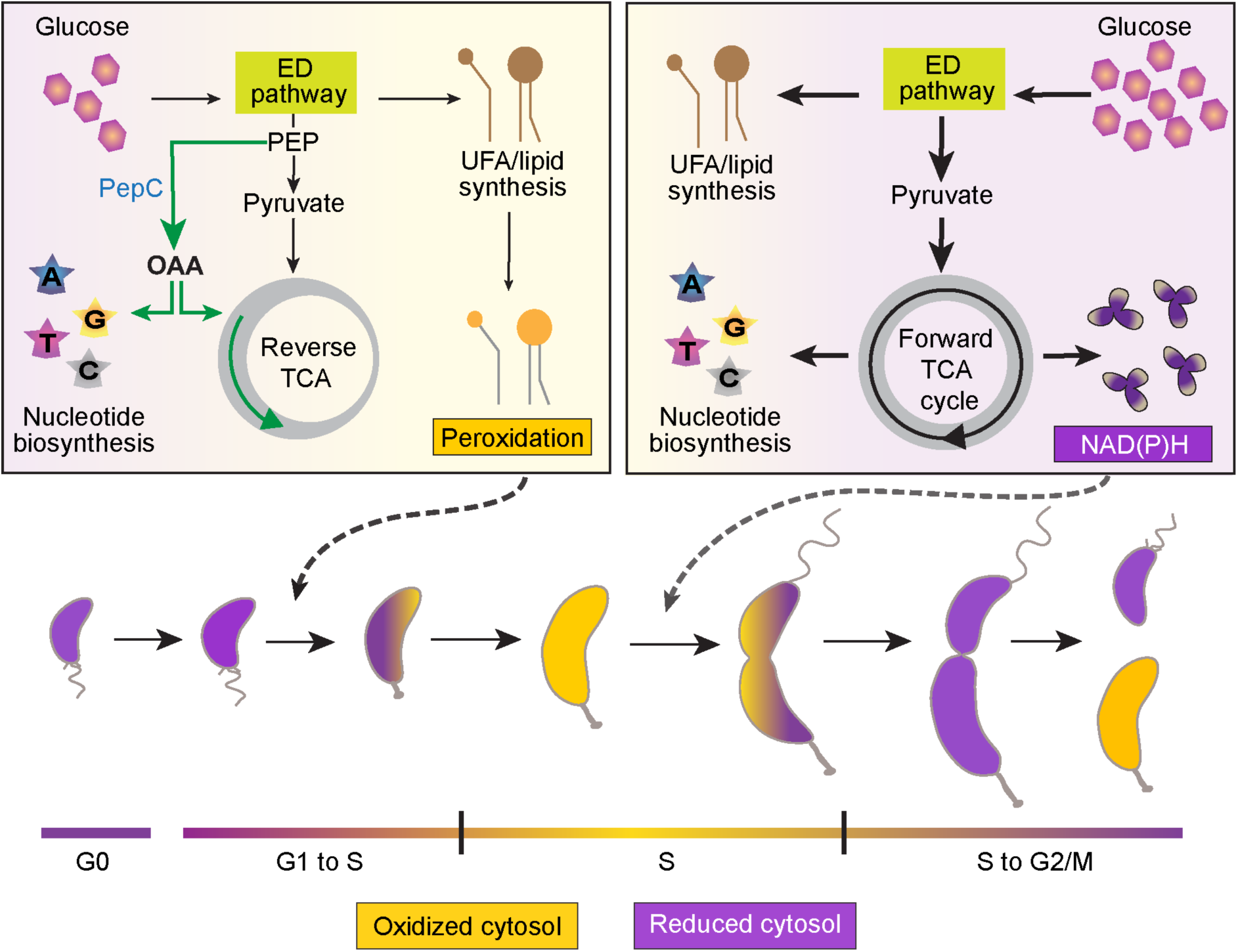
Model representing the influence of metabolism-driven redox on cell cycle transition. At the beginning of the cell cycle, glucose uptake leads to synthesis of unsaturated fatty acids (UFA) and enhancement of reverse TCA, while the forward TCA cycle is less prominent. Phosphoenolpyruvate carboxylase (PepC) catalyzed synthesis of oxaloacetate (OAA), from phosphoenolpyruvate (PEP), facilitates nucleotide synthesis. The peroxidation of UFA/lipids enables the oxidation of cytoplasm promoting G1 to S transition. At the later stages of the cell cycle, more amount of glucose uptake coupled with higher glucose metabolism and activation of forward Tricarboxylic acid (TCA) cycle enhances the production of NAD(P)H that in turn helps in reducing the cytoplasm facilitating cytokinesis. ED: Entner-Doudoroff pathway.

### Peroxidation-triggered cell cycle entry

Before initiating S-phase, wherein the genetic material is replicated, cells need to grow in size to ensure the accommodation of the newly replicating chromosome. Therefore, for proper cell cycle progression, there needs to exist a mechanism by which production of fatty acids and lipids, required for cell growth, is robustly conveyed to the cell cycle regulatory machinery. Nevertheless, our understanding of this crosstalk between metabolism-driven fatty acid synthesis and cell cycle control mechanisms have remained unclear. Our data suggest that, in *Caulobacter*, the abundant synthesis of unsaturated fatty acids, co-incident with the G1 to S transition, invariably triggers a peroxidation-dependent cytosolic oxidation. The oxidative trigger is essential for the G1 cells of *Caulobacter* to commit to S-phase. In support of this, inhibiting fatty acid synthesis prevent peroxidation and resultant cytoplasmic oxidation, thereby blocking cells from transitioning into S-phase (Figures 2D-E and H-I). Moreover, glucose catabolism, leading to enhancement of unsaturated fatty acid synthesis, a substrate for peroxidation, specifically accelerates the S-phase entry (Figure 2). Together, these results uncover lipid/fatty acid peroxidation as an unconventional signalling module that functionally couples proper metabolic progression with the triggering of S-phase entry. Notably, in *Caulobacter* once the swarmer cell transitions into a stalked cell there is a delay before the chromosome replication is initiated^41^. In light of this, we suggest that the motile swarmer cell of *Caulobacter* is in a developmentally quiescent G0 like state. The G0 to G1 entry is nutrient-dependent. We propose that the proper initiation of the G1 phase is coupled by the peroxidation driven cytosolic oxidation licensing the entry into the replicative S-phase (Figure 6). This fatty acid/lipid peroxidation-driven entry into S-phase may not unique to *Caulobacter* but could very well be prevalent in other organisms. In support of this notion, studies in breast cancer cells have shown that an overabundance of antioxidants delays S-phase entry^42^. Furthermore, this delay is accompanied by a reduction in phospholipid hydroperoxides, a key product of fatty acid/lipid peroxidation^42^. Collectively, these observations underscore the previously unrecognized physiological significance of lipid peroxidation.

We speculate that if kept within the threshold, fatty acid/lipid peroxidation can contribute to cytoplasmic oxidation, which could act as a regulatory layer during cell signalling^43^. Additionally, this redox-dependent process could aid in activating the developmental regulators that are required to facilitate the S-phase events. For example, the topoisomerase IV inhibitor NstA, which is produced co-incident with the S-phase entry^9^. The activation of NstA requires oxidation-dependent intermolecular cysteine disulphide-based dimerization to inhibit topoisomerase IV activity until late S-phase^9^. Such a type of redox-dependent regulation has also been proposed in other domains of life, albeit with no direct experimental proof^10,11^. A recent study in human retinal epithelial cells has identified a redox switch-dependent exit from cell cycle – wherein, the oxidation of a cysteine residue in the CDK inhibitor p21, decides if the cells continue to undergo proliferation^12^.

By confining peroxidation to a specific window during the cell cycle, *Caulobacter* cells ensure that there is no hyper oxidation of the cytosol that could lead to damage of the cellular components. Moreover, our data indicate that peroxidation is temporally reduced during the mid-S-phase (Figure 2G). This differential peroxidation levels could be regulated by the levels of the major antioxidant defence nucleotides, NAD(P)(H). Remarkably, the levels of NAD(P)(H), glutathione and thioredoxin are more abundant from mid-S-phase onwards (Figures 3E, 5D, and Supplemental Figures S3A, B, S5D)^44,45^, co-incident with the suppression of fatty acid/lipid peroxidation (Figure 2G). Strikingly, the TCA cycle, a major contributor towards the NADH pool, is not enhanced until mid-S-phase (Figures 3F-H, Supplemental Figure S3C and Supplemental Dataset 4). Therefore, we propose that an enhanced TCA cycle at the late-S-phase facilitates cell cycle progression by providing the reducing potential to the cell. Consistent with this model, interfering with the forward TCA cycle specifically blocks cell cycle progression at the S to G2/M transition (Figures 4A, B). Other metabolic reactions could also contribute to this enhancement of NADH pool during the late S-phase. For instance, when glutamate is used as a carbon source, the conversion of glutamate into α-ketoglutarate with concomitant reduction of NAD^+^ to NADH catalysed by the NAD-dependent glutamate dehydrogenase GdhZ is restricted to late S-phase by the cell cycle-dependent proteolytic degradation of GdhZ^7^. In addition to providing a reducing potential, this temporal increase in NADH levels likely accelerates cell division through KidO, the NADH-dependent cell division regulator^6^.

The upregulation of the TCA cycle is accompanied by increase in glucose metabolism from the mid-S-phase onwards (Figures 3F-H, Supplemental Figure S3C and Supplemental Dataset 4). Moreover, increased glucose flux may also contribute towards the increase in NADPH levels. Notably, in *Caulobacter*, NADPH is generated primarily through the Zwf-catalysed conversion of glucose-6-phosphate to 6-phospho-8-gluconolactone. NADPH produced early in the cell cycle is likely consumed by fatty acid and nucleotide synthesis, helping maintain low levels of NADPH (Figure 5D and Supplemental Figure S5D). These low levels of NADPH will promote a more oxidised cytoplasm, which in-turn supports S-phase entry. Additionally, silencing of the forward TCA cycle might contribute to carbon conservation during the early stages of cell cycle. A silent forward TCA cycle avoids the loss of carbon through decarboxylation reactions, hence saving it for fatty acid biosynthesis during the G1 to S transition.

### Temporal modulation of forward and reverse TCA facilitates cell cycle progression

The TCA cycle intermediates play a crucial role in the synthesis of amino acids that indirectly influence proper purine/pyrimidine nucleotide biosynthesis. For example, aspartate synthesised from oxaloacetate is utilized during purine and pyrimidine biosynthesis which is required for proper replication of the genetic material that happens during the S-phase^38,46^. Therefore, it is imperative that cells produce these late TCA cycle intermediates to maintain proper progression of S-phase upon transitioning from G1. Nevertheless, a robust forward TCA cycle is absent during the early stages of the cell cycle. To compensate for this, the cells operate an anaplerotic-reaction catalysed reverse TCA that enhances oxaloacetate and production of nucleotides during the early stages of the cell cycle. This highlights the importance of the temporal-specificity of reverse and forward TCA during the cell cycle. Moreover, by not operating a forward TCA, the cells ensure that NAD(P)H levels, which could favour a reduced environment in the cytosol, are not enhanced during the early S-phase. This cell cycle stage-specific operation of the forward and reverse TCA is likely enabled by cell cycle stage-specific expression of TCA cycle enzymes encoding genes. In support of this argument, cell cycle transcriptome data indicates that *pepC* transcripts are abundant specifically in the G1 cells and transcripts encoding the early TCA cycle enzymes are enhanced from mid-S-phase onwards^47^.

Another challenge the cells face is to orient the utilization of accessible nutrient substrates. A large amount of glucose could still trip the balance between the forward and reverse TCA even when the enzyme levels are low. Our results highlight the cells operate a biphasic glucose uptake to circumvent this issue. The first round of glucose uptake that happens during the early stages of the cell cycle is channelled towards nucleotide and fatty acid synthesis. The more prominent, second wave of glucose uptake from the mid-S-phase onwards (Figure 5F), facilitates enhancement of glucose metabolism and NAD(P)H levels (Figures 3E-H, 5E) creating a reducing environment.

We speculate that the differential glucose uptake observed across the cell cycle may be influenced by fluctuations in cyclic di-GMP (c-di-GMP) levels . Notably, c-di-GMP concentrations rise sharply during early S phase and decline from the mid–S phase onwards^48^. Previous studies have suggested that elevated c-di-GMP levels can inhibit glucose uptake^49^. Therefore, it is very much likely that the early increase in c-di-GMP levels suppresses glucose influx, whereas the second uptake wave is triggered once c-di-GMP levels fall below a critical threshold. Another likely scenario is the presence of two different glucose uptake mechanisms, each of which is operational at specific stages of the cell cycle. Nevertheless, our work demonstrates the occurrence of regulated glucose uptake in single cells which could likely be the case in other cell types and carry broader implications.

## Acknowledgements

We thank Sean Murray and Patrick Viollier for strains. Girish Deshpande and Kalika Prasad for comments on the manuscript. Microscopy, Flow Cytometry and Mass Spectrometry Facility at IISER Pune. Maulana Azad National Fellowship to HAG (201610068944) and Research Fellowship from IISER Pune AC. Financial support from the Science and Engineering Research Board (SERB), Department of Science and Technology, Government of India (Grant: SB/SJF/2021-22/01 to SSK), an EMBO Young Investigator Award (to SSK) and an EMBO long term fellowship (ALTF 152-2008 to RH). Research Credit from the F.R.S. – FNRS (CDR J.0169.16 to RH). RH is a Senior Research Associate of F.R.S. – FNRS. This work was supported by funds from the DBT-Wellcome Trust India Alliance through a Senior Fellowship to SKR (IA/S/20/2/505202).

## Materials and Methods

### Bacterial Strains and Medium

NA1000 and CB15 strains of *Caulobacter crescentus*, were grown on rich PYE medium (0.2% peptone, 0.1% yeast extract, 1 mM MgSO_4_, 0.5 mM CaCl_2_) or minimal (M2) medium (0.87 gl^-1^ Na_2_HPO_4_, 0.53 gl^-1^ KH_2_PO_4_, 0.25 gl^-1^ NH_4_Cl) supplemented with 0.5 mM MgSO_4_, 10 µM FeSO_4_.EDTA, 0.5 mM CaCl_2_) and incubated at 29°C unless specifically mentioned. Glucose (0.2%) was used as carbon source in M2 medium, other carbon sources, sodium aspartate (0.2%), sodium citrate (0.05%) and xylose (0.2%) were used wherever mentioned. The *Caulobacter* strains were subjected to synchronization and electroporation as previously described ^6,50^. Wherever required, kanamycin 20 µg/mL or 5 µg/mL for solid or liquid medium, respectively, Gentamycin 1 µg/mL for liquid medium and agarose (1.0%) were used. The *Escherichia coli*, EC100 (Biosearch Technologies) was used for cloning purposes and cultured in Luria Bertani (LB) medium. The culture was incubated in a rotatory shaker at 210 rpm at 37°C overnight till it reached the required OD_600_. Wherever required, kanamycin (20 µg/mL) and agar (1.25%) were used.

List of strains and plasmids used in this study are listed in Supplemental Table S3 and S4, respectively.

### Microscopy

Phase contrast microscopy was performed using Olympus IX83 inverted microscope (Evident Scientific, Japan) equipped with a U Plan X Extended Apo 100X (1.45 numerical aperture) objective and an ORCA-Flash4.0 V3 sCMOS camera (Hamamatsu, Japan). Cells were spotted on a 1% agarose pad for imaging. The acquired images were processed using ImageJ.

### Cell synchronization

The overnight *Caulobacter* (NA1000) culture, grown in M2G, (OD_600_ 0.4-0.6) was pelleted at 5000 rpm for 5 minutes at 4°C. Pellets were then washed thrice in 20 mL (for 50 mL of pelleted culture) of ice-cold 1X-M2 salt and centrifuged at 8500 rpm for 3 minutes at 4°C. The cells were then resuspended in 1.4 mL of ice-cold 1X-M2 solution without glucose. Further, 700 μl of cells were aliquoted into two 2 mL microcentrifuge tubes (MCTs). To this, 800 μl of cold Percoll (Cytiva) was added to both MCTs and mixed gently. The cell suspension was then centrifuged at 10000 rpm for 20 minutes at 4°C. Two distinct layers of cells were formed after the samples were centrifuged. The upper layer, containing the stalked cells, was carefully removed and discarded. The lower layer, consisting of swarmer cells, was resuspended in 1 mL cold 1X-M2 solution by gentle pipetting. The suspension was then centrifuged at 8500 rpm for 5 minutes at 4°C. The supernatant was removed carefully, and the cell pellet was washed thrice with cold 1X-M2 solution. Finally, the cells were resuspended in the desired growth medium and incubated at 29°C to initiate the cell cycle.

### Monitoring intracellular redox

Secondary culture (100 mL) of NA1000 *Caulobacter* cells harbouring the pSKR478-*rogfp2* plasmid was grown overnight in M2G with gentamycin in the presence of 1 mM IPTG until the cells reached an 0.4-0.6 OD_600_. The cells were then synchronized and the swarmer cell population was resuspended to an initial density of 0.4 OD_600_ in 5 mL of M2G supplemented with 1mM IPTG. A 100 µl aliquot of culture was taken at every 10-minute intervals until 140 minutes. The fluorescence intensities at 405 nm and 480 nm excitations and emission at 520 nm were measured at each 10-minute interval using a CLARIOstar plate reader (BMG LabTech). The ratio of fluorescence intensity at 520 nm for 405 nm and 480 nm excitations was calculated at each timepoint to monitor the cytosolic redox state during the cell cycle ^9,22,23^.

### Flow cytometry (FACS analysis)

Flow cytometry analyses were performed as described earlier (Narayanan et al., 2015). In brief, *Caulobacter* cells were grown in desired media conditions. A 5 µl aliquot of these cells was added to 1 mL of ice-cold 70% ethanol solution and stored overnight at -20°C or until further use. For staining and analysis, 50 µl of fixed cells were pelletized and washed thrice with 1 mL of the staining buffer (10 mM Tris HCl pH 7.2, 1 mM EDTA, 50 mM Sodium Citrate + 0.01% TritonX-100). The washed cells were then resuspended in 1mL of staining buffer containing 0.1 mg/mL RNase A (Roche Life Sciences, Switzerland) and were incubated at room temperature for 1 h. The cells were then harvested by centrifugation at 10,000 rpm for 3 min. The pellets were resuspended in 100 µl of staining buffer supplemented with 0.5 mM SYTOX^TM^ Green Nucleic Acid Stain (Invitrogen) followed by incubation in dark for 5 min. These cells were then analyzed immediately on BD Accuri^TM^ C6 Flow Cytometer (BD Biosciences, San Jose, California, USA). Chromosome number was directly estimated from the green fluorescence (FL1-A) value of the stained cells and the data was analyzed using FlowJo software to determine the number of cells with one chromosome (1N) to two chromosomes (2N).

To monitor cell cycle progression in the synchronized population of WT NA1000, swarmer cells were resuspended to an initial density of 1.2 OD_600_ in the respective media conditions. Samples were then collected at every 20 minutes during the cell cycle and processed as mentioned above.

### Genetic screen

The genetic screen was described in ^13^. Briefly, a population of CB15 Δ*pilA* (UJ590) was maintained in exponential growth phase in M2G for more than 2,000 generations by diluting growing cells every 24 hrs. Aliquots of the evolved population were regularly stored at -80°C. Genomic DNA were extracted from the ancestor (UJ590) and the population evolved for more than 2,000 generations (UJ4825), and used for Illumina sequencing (GA-II, Fasteris).

### qPCR analysis

RNA extraction was performed using RNeasy mini kit (QIAGEN) according to the manufacturer’s instructions. Then cDNA was synthesized by using the High Capacity cDNA Reverse Transcription Kit (Applied Biosystems) according to the manufacturer’s instructions. Briefly, RNA (∼0.5 µg) was added to Reverse Transcription Master Mix containing RT buffer, dNTPs (1mM each), random primers and MultiScribe Reverse Transcriptase (50 U) in a 20 µl final volume reaction. The reaction was incubated at 25°C for 10 mins, then at 37°C for 2 hrs and finally at 85°C for 5 mins. RT-qPCR was performed using FastStart Universal SYBR Green Master (Roche) according to the manufacturer’s instructions. Briefly, 5 µl of cDNA was added to FastStart Universal SYBR Green Master Mix and pairs of oligonucleotides for each gene of the *zwf* operon or for 16S used as a normalizer (568/569 for *zwf*, 570/571 for *pgl*, 572/573 for *edd*, 574/575 for *glk*, 558/559 for 16S) in a 20 µl final volume reaction. The reactions were then performed at least in triplicate in a LightCycler 96 Instrument (Roche). For a given strain grown in a given condition, ΔC_T_ values were calculated by subtracting C_T_ mean values of 16S to the C_T_ mean values of each targeted gene. Then, for each gene, ΔΔC_T_ values were calculated by subtracting ΔC_T_ values of the reference strain to the ΔC_T_ values of a mutant strain. Finally, data were expressed as log_2_[FC] of ΔΔC_T_ values.

### Promoter activity assay

The β-galactosidase-based promoter activity assay was done as mentioned previously (Narayanan et al., 2015). In brief, the overnight cultures harboring the *lacZ* reporter were sub-cultured and grown at 29°C till the cells reached 0.1-0.4 OD_660_ (A660). To 50 μL of the cells, 10 μL of chloroform was added, followed by the addition of 750 μl of Z-buffer (60 mM Na_2_HPO_4_, 40 mM NaH_2_PO_4_, 10 mM KCl, 1 mM MgSO_4_.7H2O, pH 7.0). Next, 200 µl of ortho-Nitrophenyl-b-D-galactoside (4 mg/mL ONPG, dissolved in 0.1M potassium phosphate buffer [pH 7.0]) was added in above mixture. The reaction mixture was incubated at 30°C on thermo-block till a yellow colour was developed and the incubation time was noted (t). Finally, 500 µl of 1 M Na_2_CO_3_ solution was added to stop the reaction and the absorbance at 420 nm (A420) of the supernatant was monitored, where Z-buffer was used as the blank. The miller units (U) were calculated using the equation U= (A420nm X 1000) / (A660nm X t X v), where ‘t’ is the incubation time (min), ‘v’ is the volume of culture taken (µl).

### Enzyme activity assay

Total proteins were extracted from cultures (10 - 50 ml) grown until mid-exponential phase (OD_660_ ∼0.3). Then cells were pelleted at 6,000 g for 10’ at 4°C and washed once in 10 ml of 20 mM phosphate buffer (12.5 mM Na_2_HPO_4_, 7.5 mM KH_2_PO_4_). The pellet was resuspended in 2 ml extraction buffer (40 mM HEPES buffer pH 7.4, 10 mM MgCl_2_, 1 mM EDTA, 1 mM EGTA, 1 mM benzamidine, 1 mM amino n-capronic acid, 20 µM leupeptin, 250 µM DTT, 1mM PMSF, 2% glycerol) and sonicated 6x for 10 secs. Cell debris were pelleted 15,000 g for 15 min at 4°C and discarded. Proteins extracts were quantified using Bradford assay. Glucose-6-phosphate dehydrogenase (Zwf) and Glucokinase (Glk) activities were determined by measuring NADP^+^ reduction to NADPH at 340 nm at 25° C until the slope stabilises. For both reactions, proteins (∼2 µg) were added to assay buffer containing 100 mM Tricine/KOH pH 8.0, 10 mM MgCl_2_ and 0.05 % Triton X100. For Zwf assays, 40 mM NADP^+^ and 50 mM Glucose-6-Phosphate were added to the mixture. For Glk assays, 40 mM NADP^+^, 200 mM Glucose, 2 U of Zwf grade II (Roche) and 8 mM ATP were added to the mixture. The reactions were performed 25°C in a 40 µl final volume reaction in a plate reader (ELx808, Biotek) and analysed with Gen5 Data Analysis Software (Biotek).

### Growth curve and dilution spotting

Overnight cultures of *Caulobacter* cells, grown in desired conditions, were washed twice with sterile MilliQ (MQ) water and sub-cultured in desired growth media with an initial OD_600_ of 0.025 or 0.1 for growth curve and dilution spotting, respectively. For growth curve experiments, 200 µl of the above 0.025 OD_600_ cells was taken in a 96-well clear-bottom transparent plate. Plates were placed in a CLARIOstar plate reader (BMG LabTech) at 29°C with continuous double-orbital shaking at 500 rpm. OD_600_ was recorded every 30 minutes. For dilution spotting, 10 µl of serially diluted culture was spotted on Petri plates containing 1% agar with respective growth media. The plates were incubated at 29°C and imaged after 48 h using Amersham ImageQuant 800 Imager (Cytiva, USA).

### Metabolite extraction

*Caulobacter* cells were grown in the desired media and conditions as mentioned. Cells were quenched by pelleting 1 mL of 0.8 OD_600_ cells at 14,000 rpm for 3 min at 4°C and immediately resuspended into 2 mL of chilled 60% methanol containing 5 mM Tricine buffer (pH 7.4). The quenched cells were pelleted at 14,000 rpm for 3 min at 4°C and the supernatant was discarded. The pellet was flash-frozen in liquid nitrogen and immediately stored at -80°C until the extraction of lipids or metabolites.

Polar metabolites were extracted using the protocol as described earlier ^51^. Briefly, the frozen cell pellets were thawed on ice for 5 min resuspended in 600 μl of 75% (v/v) ethanol. The ethanol had respective internal standards: 5-10 nmol of either D-glucose-1-^13^C (Merck, catalog # 297046) or U-^13^C_6_-Glucose (Cambridge Isotope Laboratories, catalog # CLM-1396-10) for non-derivatized metabolites, and 0.5-1 nmol of D4-succinic acid (Cambridge Isotope Laboratories, catalog# DLM-2307) for derivatized metabolites. The cells were then incubated at 80°C for 3 min on a thermal block, with constant shaking, and immediately plunged into ice for 5 mins to enable lysis. The lysed samples were then centrifuged at 14,000 rpm for 10 min at 4°C, to remove the cell debris, and the supernatant was transferred to a fresh MCT. The supernatant containing the extract was then dried under vacuum using CentriVap vacuum concentrator (Labconco) and stored at -80°C until LC-MS/MS analysis.

For derivatization of polar metabolites (TCA cycle intermediates), the dried extract was reconstituted in 150 µl of MilliQ water and 75 μl of 1 M N-(3 dimethylaminopropyl)-N’-ethylcarbodiimide (prepared in 13.5 mM pyridine buffer, pH 5.0) (Sigma, catalog# E7750) was added and mixed gently. Further, 150 μl of 0.5 M O-benzylhydroxyl amine (Sigma, catalog# B22984), prepared in 13.5 mM pyridine buffer at pH 5.0, was added. The above mixture was incubated at 25°C for 1 h with continuous shaking on a thermal block. To this reaction mixture 350 μl of ethyl acetate was added and mixed by shaking for 10 min on a thermal block at 25°C. The samples were then centrifuged at 3000xg for 5 min at 4°C. The upper layer was carefully transferred into a fresh MCT. The ethyl acetate-based extraction was repeated two more times, and the upper layer from all three extractions were pooled together. The pooled ethyl acetate fractions were then dried under vacuum using CentriVap vacuum concentrator (Labconco), and stored at -80°C until LC-MS/MS analysis.

Fatty acids and lipids were extracted using a previously mentioned protocol ^51^. Briefly, the frozen cell pellets were thawed on ice for 5 min and resuspended in 1 mL of ice-cold 1X PBS (8 g/litre NaCl, 0.2 g/litre KCl, 1.44 g/litre Na_2_HPO_4_ and 0.24 g/litre KH_2_PO_4_; pH 7.4). The cell suspension was transferred into a fresh glass vial and 3 mL of Chloroform:Methanol (2:1) mixture containing 1 nmol of LysoPS17:1 (Avanti Polar Lipids, catalog# 858141) as internal standard was added. The mixture was vigorously vortexed and centrifuged at 3,000 rpm for 15 min at room temperature. Two distinct layers were formed with a protein disc at the interface. The bottom organic layer was transferred into a new glass vial. To enable the extraction of lipids, that remain in aqueous layer 50 µl formic acid was added and vortexed before adding 2 mL of chloroform. The mixture was then vortexed, and centrifuged at 3,000 rpm for 15 min at room temperature. The bottom organic layer was removed and pooled up with the previously extracted sample. The pooled samples were then dried in a stream of pure nitrogen gas and stored at -80°C until LC-MS/MS analysis.

For cell cycle time course experiments, overnight grown *Caulobacter* cells of having OD_600_ of 0.4-0.6 were synchronized using Percoll as described above. The synchronised population of swarmer cells were resuspended, in M2G media containing U-^13^C_6_-Glucose (Cambridge Isotope Laboratories, catalog # CLM-1396-10) as the sole carbon source, to a final OD_600_ of 0.8. Equal volume (1 mL) from the resuspended 0.8 OD_600_ cells were collected at every 20 mins interval during the cell cycle. The samples were then quenched and processed as described above for extractions of lipids, non-derivatized and derivatized metabolites. For non-derivatized metabolites, 5-10 nmol of D-glucose-1-^13^C [Merck, catalog # 297046) was used as an internal standard.

### LC-MS/MS analysis

All LC-MS runs were carried out using the AutoMS/MS acquisition method on an Agilent 6545 Q-TOF mass spectrometer fitted with an Agilent 1290 Infinity II UHPLC system as described previously with minor modifications. The dried metabolites and fatty acids/lipids were re-solubilized in appropriate solvents. For non-derivatized metabolites, a mixture of H_2_O:Acetonitrile (19:1) and 5 mM Ammonium Acetate was used. The derivatized metabolites were rehydrated in 50 µl of MeOH:H_2_O (1:1). To resuspend the lipid extracts, 200 µl of CHCl_3_:CH_3_OH (2:1) was used. 10 μL of the resuspended samples was injected onto either a Phenomenex Synergi Fusion-RP Column (150 mm x 4.6 mm, 4 μm, 80 Å) (catalog # 00F-4424-E0) (For Polar Metabolites) or a Phenomenex Gemini C18 column (50 mm × 4.6 mm, 5 μm, 110 Å) (catalog # 00B-4435-E0) (for lipids) fitted with a Phenomenex guard column (3.2 mm X 8.0 mm) (catalog # KJ0-4282). All details of the LC profiles and MS Parameters are provided in Supplemental Tables S5 and S6.

Data analysis for metabolites (non-derivatized and derivatized) was performed using Agilent MassHunter Qualitative Analysis 10.0 software, and all the peaks were manually validated based on relative retention times and fragments obtained ^51,52^ if any. All detected species were within a mass accuracy of 15 ppm and quantified by measuring the area under the curve for different metabolites that was normalized to levels in blank (if any) and levels of respective internal standard (D-glucose-1-^13^C, U-^13^C_6_-Glucose or D4-succinic acid) and OD_600_ of the sample.

For data analysis of lipids, a library for different *Caulobacter* lipid species was curated as a Personal Compound Database Library (PCDL) using MassHunter PCDL manager B.08.00 (Agilent Technologies). This library was curated using the METLIN lipid library as reference. The data files were processed in Agilent MassHunter Qualitative Analysis 10.0 using the PCDL library, and all the peaks were validated based on relative retention times and fragments obtained ^51^ if any. All detected species were within a mass accuracy of 15 ppm and quantified by measuring the area under the curve (AUC) for different lipid species which were then normalized to levels in blank (if any) and levels of internal standard (C17:1 Lyso-PS) and OD_600_ of the sample.

### Fatty acid/Lipid peroxidation

Thiobarbituric acid responsive substances (TBARS) assay was employed to measure fatty acid/lipid peroxidation in *Caulobacter* cells. The TBARS assay was performed as mentioned previously ^53,54^ with slight modifications. Briefly, 1 mL of 1 OD_600_ cells were pelleted at 12,000 rpm for 5 mins at 4°C. The pellets were washed in 1 mL of cold 50 mM K_2_HPO_4_ buffer (pH 7.4) containing 0.1 mM of butylated hydroxytoluene (BHT, dissolved in ethanol) and 1 mM of phenylmethanesulfonyl fluoride (PMSF) dissolved in ethanol. The washed pellet was then resuspended in the 50 mM K_2_HPO_4_ buffer and lysed by sonication in cold conditions. The lysed cells were centrifuged at 14,000 rpm for 15 min at 4°C. To 900 µl of the supernatant, 250 µl of chilled 100% trichloroacetic acid (TCA, dissolved in distilled water) was added. Samples were mixed by pipetting and kept on ice for 10 mins, and centrifuged at 12,000 rpm for 5 mins at 4°C. To 1 mL of the supernatant, 2 mL of freshly prepared 0.3% Thiobarbituric acid solution (prepared in 0.1 M HCl) containing 10 mM of BHT was added. The reaction mixture was incubated at 100°C for 1 h and then cooled to room temperature (RT) for 20 mins. 500 µl of the reaction mixture was transferred to fresh MCT to which equal volume of butanol was added. Samples were centrifuged at 4,000xg for 10 mins at RT. After centrifugation, 100 µL of the upper layer was collected and the fluorescence intensity was measured at an excitation of 515 nm and an emission of 555 nm using a Tecan microplate reader (Tecan, Switzerland). TBARS binds to malondialdehyde (MDA), a by-product of lipid peroxidation, forming a complex which gives fluorescence at 515 nm excitation.

### Depletion experiments

For the depletion of isocitrate dehydrogenase (*idH)*, the *Caulobacter idh*::*P_van_-idh* cells were grown in PYE supplemented with 0.5 mM vanillate. The cells were pelleted and washed three times using PYE to remove vanillate. The washed cells were resuspended to 0.2 OD_600_ into PYE media without vanillate. The cells were then incubated for 7 h at 29°C to allow the depletion of IDH. The depleted cells were then used for microscopy and flow cytometry analysis. For mass spectrometry, cells equivalent to 1 mL of 0.8 OD_600_ were quenched as described below.

Depletion of *fabH* was carried out as previously mentioned ^18^. Briefly, Δ*fabH vanA::*P*_van_-fabH* cells grown overnight in M2G supplemented with vanillate was washed thrice with 1X M2 salt solution to remove the vanillate. The cells were then resuspended in M2G media without vanillate to 0.2 OD_600_. The cells were incubated for 7 h at 29°C and sub-cultured, into M2G with or without vanillate for 16 h and used for analyses using flow cytometry. For fatty acid supplemented growth analyses of *fabH*-depleted cells, the cells depleted for 16 h were sub-cultured into M2G medium supplemented with 1% tergitol or 100 μM sodium oleate or 100 μM sodium stearate, and the growth of these cells were monitored as described above. For flow cytometry analysis, lipid peroxidation and cytosolic redox measurements, *fabH* depleted cells were grown in M2G media containing either 1% tergitol or 100 μM sodium oleate for 16 h. The cells were then synchronized and resuspended in the respective media and samples collected at desired time intervals were used for analyses. Note that sodium oleate and sodium stearate were dissolved in the presence of 1% tergitol.

#### NAD(P)H/NAD(P)^+^ measurement-Colorimetric assay

The extraction and analyses of NAD^+^ and NADH were carried out by using NAD^+^/NADH quantification kit (Sigma: MAK037) as per the manufacturer’s protocol, with slight modifications. Briefly, 1 mL of 0.8 OD_600_ cells was pelleted at 8,500 rpm for 3 min at 4°C and resuspended in 500 µl of ice-cold 1X PBS (8 g/L NaCl, 0.2 g/L KCl, 1.44 g/L Na_2_HPO_4_ and 0.24 g/L KH_2_PO_4_; pH 7.4). The resuspended cells were then sonicated in cold-conditions. The lysate was centrifuged at 14,000 rpm for 5 min at 4°C to remove cell debris. Two aliquots of 200 µl supernatant were used for further analyses. One of the aliquots was heated at 60°C for 30 min to specifically decompose NAD^+^. The other aliquot, frozen on dry ice, was used to measure total NAD(H) levels. For NAD(H) quantification, 50 µl sample, 98 µl cycling buffer, and 2 µl cycling enzyme mix were mixed together and incubated at RT for 10 min. To this, 10 µl of NADH developer was added and kept in dark. After 2 h of incubation, absorbance was measured at 450 nm using CLARIOstar plate reader (BMG LabTech). Absorbance at 450 nm (A450nm) was normalized to blank (reaction mixture with 1XPBS). NAD^+^ was quantified by measuring the difference in absorbance of total NAD(H) to NADH aliquots. A standard curve was generated using the NADH standard provided with the kit.

For the quantification of NADP(H), the NADP^+^/NADPH quantification kit (Sigma: MAK038) was used following the manufacturer’s protocol, similar to that described above for NAD(H).

#### NAD(P)H/NAD(P)^+^ measurement-Luminescence based assay

The extraction and analyses of NAD^+^ and NADH by luminescence-based assay were done using NAD/NADH-Glo^TM^ Assay kit (Promega: G9071) following the manufacturer’s protocol, with slight modifications. Briefly, 1 mL of 0.8 OD_600_ cells was pelleted at 8,500 rpm for 3 min at 4°C and resuspended in 150 µl of ice-cold 1X PBS buffer (8 g/L NaCl, 0.2 g/L KCl, 1.44 g/L Na_2_HPO_4_ and 0.24 g/L KH_2_PO_4_; pH 7.4) and 150 µl of 0.2N NaOH. Cells were sonicated in cold conditions and cell debris were removed by centrifugation at 14,000 rpm for 5 min at 4°C. 100 ul supernatant was aliquoted into two MCTs. To facilitate NAD^+^ extraction, 50 µl of 0.4N HCl was added to one of the aliquots. Both aliquots were then heated at 60°C for 15 min and then cooled to RT for 10 mins. The aliquot for NAD^+^ extraction, was mixed with 50 µl of 0.5M Tris-base (pH 10) while the other aliquot (for NADH extraction), was mixed with 100 µl of 1M Tris-HCl (pH 8.0). For quantification of NAD(H), a reaction mix consisting of 25 µl of the extracted sample was mixed with 25 µl of NAD/NADH Glo Detection Reagent. The reaction mix was incubated at RT in dark for 1 h and luminescence was measured using CLARIOstar plate reader (BMG LabTech). Luminescence of samples were normalized with blank (25 µl 1XPBS with 25 µl NAD/NADH Glo detection reagent). A standard curve was generated using NADH standard to determine the absolute amount of NADH or NAD^+^ in the samples.

For the quantification of NADP(H), the NADP^+^/NADPH-Glo^TM^ Assay kit (Promega: G9081) was used following the manufacturer’s protocol, similar to that described above for NAD(H).

#### NADH/NAD^+^ measurement-Peredox-mCherry genetic sensor

We employed Peredox_mCherry (pRsetB-His7tag-Peredox-mCherry) to quantify the cytosolic NADH/NAD^+^ ratio ^55^. Peredox_mCherry consists of two components: (i) an NADH binding protein T-Rex and (ii) a pH-resistant T-Sapphire protein (cpFP,) producing green fluorescence, which is linked between the T-Rex dimer. T-Sapphire gives the readout of cytoplasmic NADH/NAD+ ratio due to the binding competition between NADH and NAD^+^ with T-Rex protein. Binding of NAD+ gives an open conformation whereas binding of NADH gives a closed confirmation leading to increased fluorescence from T-sapphire^55^. The probe is linked with mCherry (red fluorescence) as a control for the protein copy number and mCherry fluoresces irrespective of the NADH binding.

*Wild-type Caulobacter* cells producing peredeox-mCherry from the vanillate inducible promoter at the chromosome (*vanA*::P*_van_*-*perdox-mCherry*) was synchronised after growing in the presence of 0.5 mM vanillate for 3 h to induce the production of peredox-mCherry. The synchronised swarmer cells were resuspended in M2G supplemented with 0.5 mM vanillate. For NAD(P)(H) quantification, 100 µl of cells were collected at at regular intervals of 20 min and pelleted at 9,000 rpm for 3 min. The pellets were resuspended using 1 mL of 1X M2 salt. 100 µl of the resuspended cells were added into 96 well black bottom plate and the fluorescence was measured using CLARIOstar plate reader (BMG LabTech). The Peredox fluorescence was measured using excitation 405 nm and emission 510 nm. The mCherry intensity was measured using excitation 587 nm and emission 625 nm. The NADH/NAD+ was determined by taking the ratio of fluorescence from Peredox to mCherry after normalizing with the intensities at their respective blanks in 1X M2 salt.

### 2-NBD Glucose uptake assay

Glucose uptake in the *Caulobacter* cells was determined by using a non-metabolizable, fluorescent analog of glucose, i.e., 2-NBDG (2-Deoxy-2-[(7-nitro-2,1,3-benzoxadiazol-4-yl) amino]-D-glucose) as described earlier ^56,57^. Briefly, overnight culture of *Caulobacter* was synchronized and cells were resuspended in M2G. The glucose uptake was monitored at every 20 min during the cell cycle. At each timepoint, 100 µM of 2-NBDG was added into 100 µl of 1.0 OD_600_ cells. Cells were incubated in dark for 2 min at 29°C and then immediately centrifuged at 12,000 rpm for 3 min. The cell pellet was then washed with 1 mL of 1XPBS. The washed cells were resuspended in 100 µl of 1X PBS, and fluorescence was monitored using CLARIOstar plate reader (BMG LabTech) in black 96-well Nunc plates using excitation at 465 nm and emission at 540nm. As control, at each time point, cells without 2NBDG were processed and monitored simultaneously.

