## Supplemental Information for "A metabolic hierarchy directs cell cycle transition and morphogenesis"

### SUPPLEMENTAL FIGURE

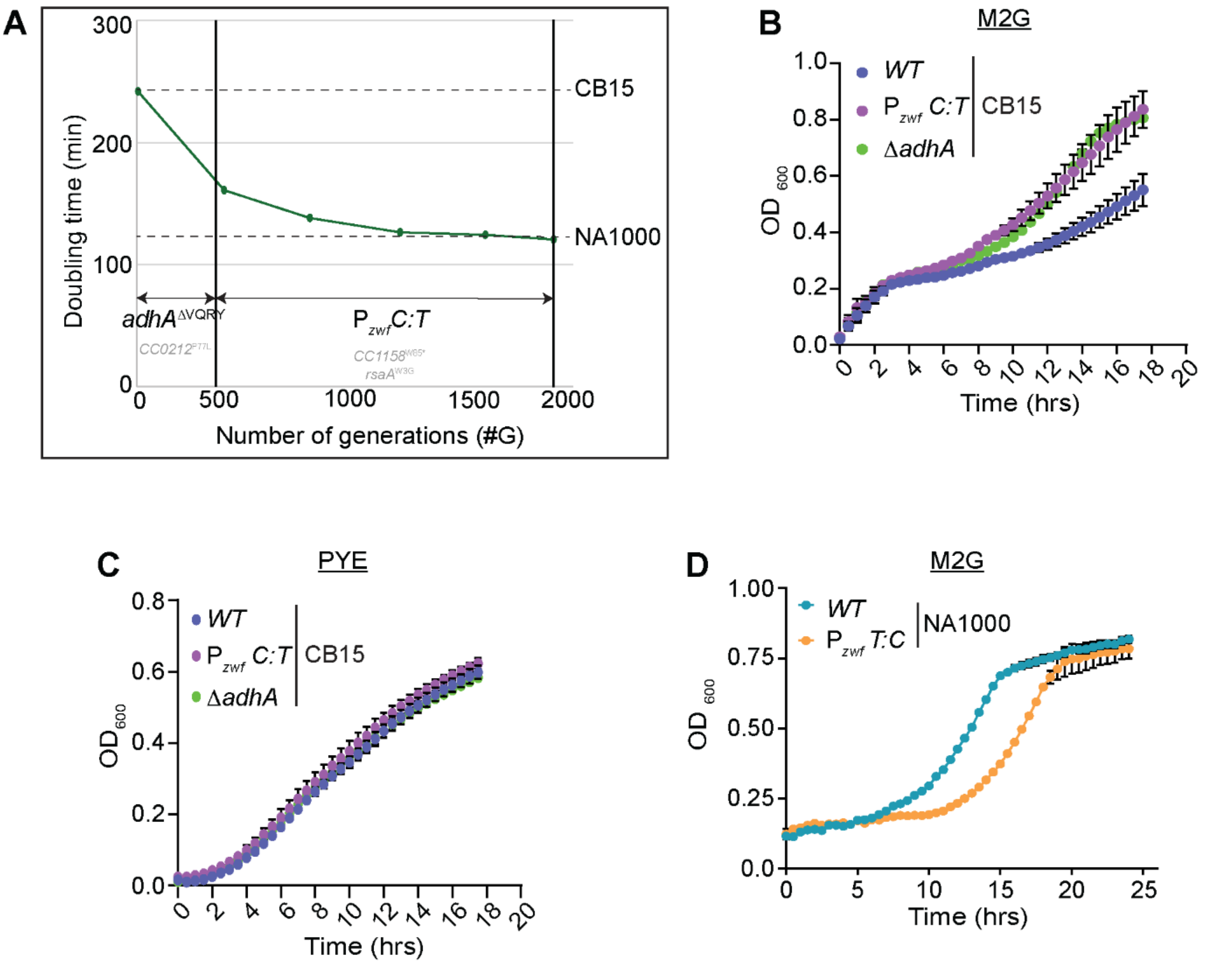


**Supplemental Figure S1: An experimental evolution screen identified fast-growing mutants targeting genes related to metabolism. Related to Figure 1. (A)** Selection of mutations that progressively decreased the doubling time of the ancestor (CB15 Δ*pilA*) growth for 2,000 generations in M2G. Each point corresponds to the doubling time of intermediate timepoints stored during the evolution experiment. Two mutations were selected during the first 500 generations (*adhA*$^{\Delta VQRY}$ and *CC0212*$^{P77L}$) and three additional mutations were selected in during the next 1,500 generations ($P_{zwf}$*C:T*, *CC1158*$^{W85*}$ and *rsaA*$^{W3G}$). (**B-C**) Graph representing the growth of *WT* CB15 and CB15 cells harbouring the mutations $P_{zwf}$*C:T* or Δ*adhA* in **(B)** M2G and **(C)** PYE. **(D)** Graph representing the growth of *WT* NA1000 and NA1000 harbouring the $P_{zwf}$*T:C* mutation in M2G. The data represented in B-D are from three independent biological replicates, ±SE.

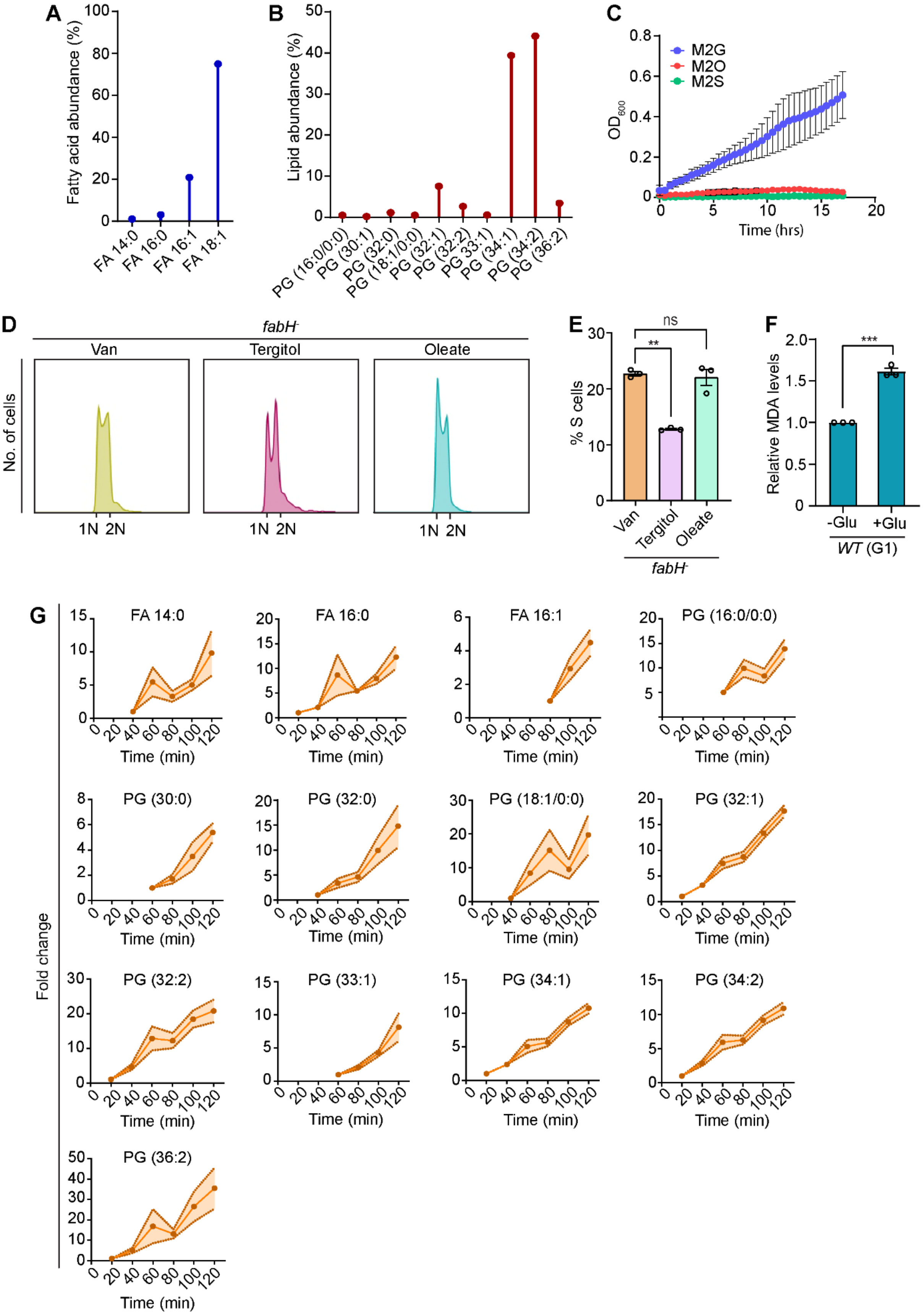
A
Fatty acid abundance (%)
FA 14:0
FA 16:0
FA 16:1
FA 18:1
B
Lipid abundance (%)
PG (16:0/0:0)
PG (30:1)
PG (32:0)
PG (18:1/0:0)
PG (32:1)
PG (32:2)
PG 33:1)
PG (34:1)
PG (34:2)
PG (36:2)
C
M2G
M2O
M2S
OD600
Time (hrs)
D
fabH-
Van
Tergitol
Oleate
No. of cells
1N 2N
E
ns
**
% S cells
Van
Tergitol
Oleate
fabH-
F
***
Relative MDA levels
-Glu
+Glu
WT (G1)
G
Fold change
FA 14:0
FA 16:0
FA 16:1
PG (16:0/0:0)
PG (30:0)
PG (32:0)
PG (18:1/0:0)
PG (32:1)
PG (32:2)
PG (33:1)
PG (34:1)
PG (34:2)
PG (36:2)
Time (min)

**Supplemental Figure S2: $^{13}$C-glucose based flux analysis showed high abundance of unsaturated fatty acid/lipid, synthesized during the cell cycle. Related to Figure 2.** Bar graph representing the % abundance of $^{13}$C-derived various species of **(A)** fatty acids (FA) and **(B)** phosphatidylglycerol (PG) in *WT* NA1000 cells. The data obtained from mass spectrometry, derived from eleven independent biological replicates. **(C)** Graph representing the growth of *WT* NA1000 in M2 medium supplemented either with glucose (M2G), 100 µM oleate (M2O) or 100 µM stearate (M2S). The data represented are from three independent biological replicates, ±SE.
(D) Flow cytometry profiles, representing 1N and 2N DNA content and (E) Bar graph representing % S-phase population, in *fabH* depleted (*fabH*$^{-}$) cells grown in M2G supplemented with 100 µM oleate dissolved in 1% tergitol. Cells grown in the presence of 1% tergitol was used as control. (F) Fatty acid/lipid peroxidation represented as relative malonaldehyde (MDA) levels in G1 cells resuspended in M2 medium with or without glucose. Synchronised population of glucose depleted *WT* NA1000 cells were used. Data represented in E and F are from three independent biological experiments, ±SE. (G) Mass spectrometry-based quantification of $^{13}C_6$-glucose derived fatty acid (FA) and phosphatidyl glycerol (PG) species during the cell cycle in *WT* NA1000 cells. Cells were synchronized and analyzed at every 20 minutes during the cell cycle in M2G medium containing $^{13}C_6$-glucose as the sole carbon source. The data is derived from eleven independent biological replicates, ±SE, and is relative to the initial time-point, where each species were detected. Statistical analyses were done using an ordinary one-way ANOVA with Dunnett's multiple comparisons test in E; and an unpaired two-tailed t-test in F ***$p$ = 0.0002; **$p$ = 0.0021, ns = not significant.

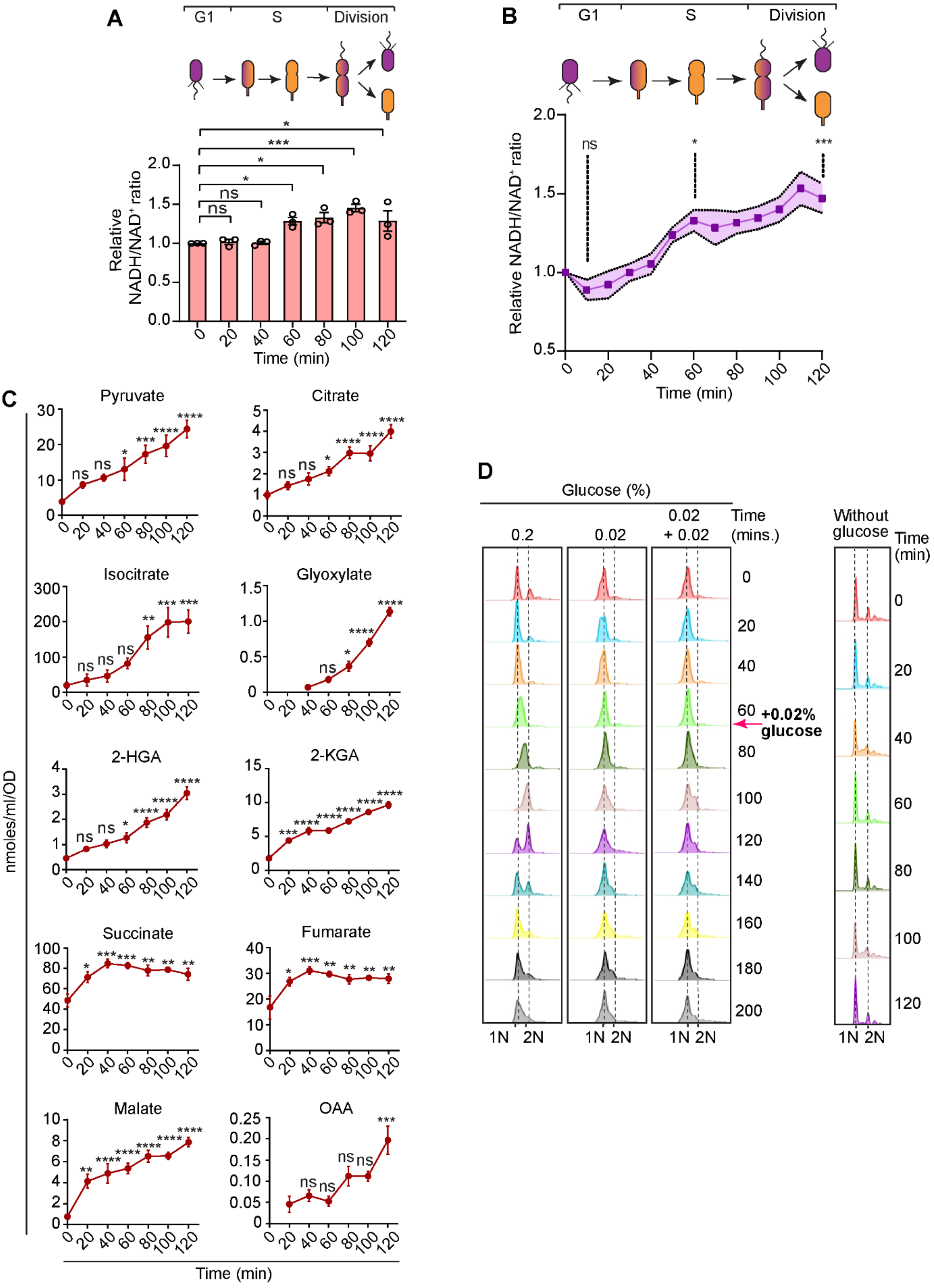


**Supplemental Figure S3: Forward TCA cycle is enhanced from mid-S-phase onwards. Related to Figure 3.** NADH/NAD$^+$ ratio during the cell cycle in synchronized population of *WT* NA1000 cells. The experiment was performed using

**(A)** luminescence-based assay and **(B)** NADH/$NAD^+$ sensing by Peredox mCherry genetic sensor. Data represented are from three independent biological replicates, ±SE, and relative to the abundance at 0 min. **(C)** Mass spectrometry-based quantification of $^{13}C_6$-glucose derived TCA cycle metabolites during the cell cycle in *WT* cells. Cells were synchronized and analyzed at every 20 minutes during the cell cycle in M2G medium containing $^{13}C_6$-glucose as the sole carbon source. The data is derived from at least five independent biological replicates, ±SE, and is relative to the initial time-point (0 min). **(D)** Flow cytometry profiles, representing 1N and 2N DNA content, during the cell cycle in synchronized population of *WT* NA1000 cells grown in M2 medium with 0.2% or 0.02% or 0.02% + 0.02% glucose (additional 0.02% glucose added at 60 min during cell cycle) or without glucose. (*Abbreviations: OAA: oxaloacetic acid, 2-KGA: α-ketoglutaric acid, 2-HGA: hydroxyglutaric acid*.) Statistical analyses were done using an ordinary one-way ANOVA with Dunnett's multiple comparisons test in A-C; ****p < 0.0001, ***p = 0.0002; **p = 0.0021, *p = 0.0332, ns = not significant.

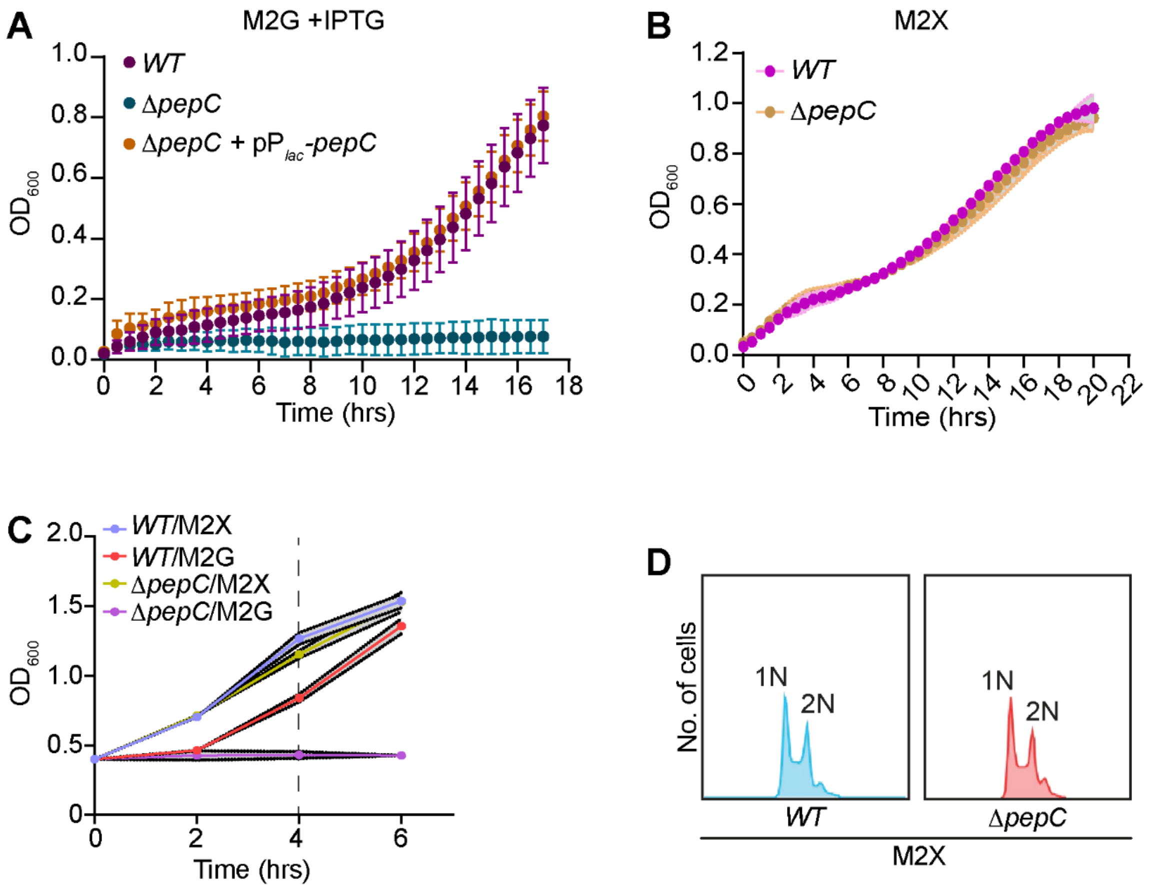


**Supplemental Figure S4: Phosphoenolpyruvate carboxylase (PepC) activity facilitates reverse TCA during early S-phase. Related to Figure 4. (A)** Graph representing the growth of *WT*, *pepC* null (Δ*pepC*) and Δ*pepC* cells ectopically expressing *pepC* from a $P_{lac}$ promoter on a medium copy vector (Δ*pepC* + $pP_{lac}$-*pepC*) in M2G medium with 1mM IPTG. **(B)** Graph representing the growth of *WT* and Δ*pepC* in M2 medium with xylose (M2X). **(C)** Growth of *WT* and Δ*pepC* in M2G and M2X medium. **(D)** Flow cytometry profiles of *WT* and Δ*pepC* cells grown in M2X for 4h. For C and D cells grown overnight in M2X were used. The cells were washed and resuspended either in M2G or M2X.

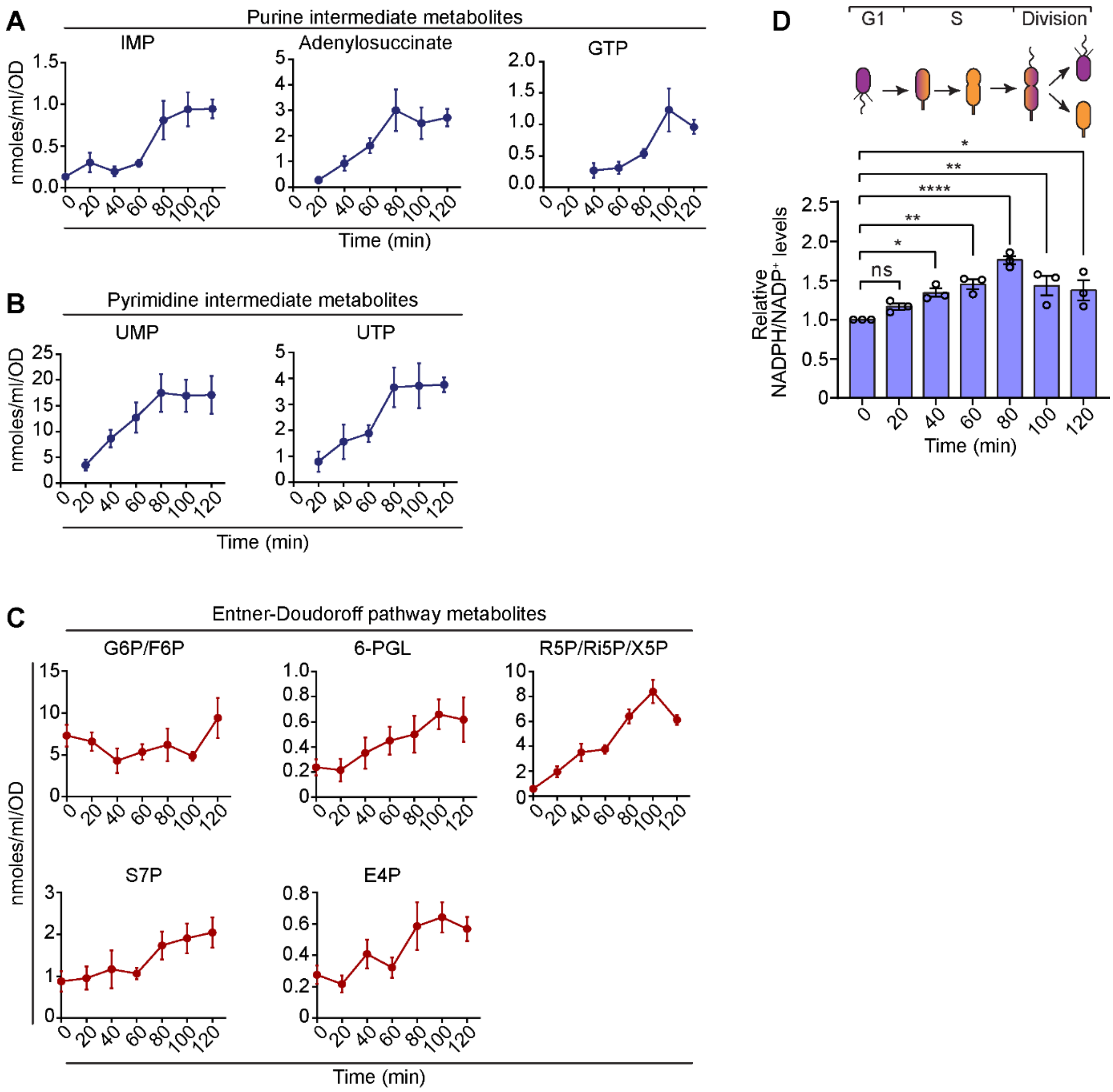


**Supplemental Figure S5: Nucleotide biosynthesis during the cell cycle in *Caulobacter crescentus*. Related to Figure 5.** Mass spectrometry-based quantification of $^{13}C_6$-glucose derived (A) purine intermediates (B) pyrimidine intermediates and (C) Entner-Doudoroff pathway intermediate metabolites during the cell cycle in *WT* cells. Cells were synchronized and analyzed at every 20 minutes during the cell cycle in M2G medium containing $^{13}C_6$-glucose as the sole carbon source. The data is derived from five independent biological replicates, ±SE, and is relative to the initial time-point (0 min). (D) $NADPH/NADP^+$ ratio during the cell cycle in synchronized population of *WT* NA1000 cells. The experiment was performed using luminescence-based assay. Data represented are from three independent biological replicates, ±SE, and relative to the abundance at 0 min. Statistical analyses were done using an ordinary one-way ANOVA with Dunnett's multiple comparisons test in D; ****p

< 0.0001; **p = 0.0021, *p = 0.0332, ns = not significant. (*Abbreviations: IMP: inosine monophosphate, GTP: guanosine triphosphate, UMP: uridine monophosphate, UTP: uridine triphosphate, G6P: glucose-6-phosphate, F6P: fructose-6-phosphate, 6-PGL: 6-phosphogluconolactone, Ri5P: ribulose-5-phosphate, R5P: ribose-5-phosphate, X5P: xylose-5-phosphate, E4P: erythrose-4-phosphate, S7P: sedoheptulose-5-phosphate)*

### SUPPLEMENTAL TABLES

| Doubling Time (min) | | | | | |
|---|---|---|---|---|---|
| **CB15 ∆*pilA* background** | **M2G** | **M2X** | **PYE** | **PYEG** | **PYEX** |
| Parental | 242 ± 1 + | 166 ± 1 | 102 ± 1 | 102 ± 2 | 100 ± 1 |
| *adhA*$^{\Delta VQRY}$ | 161 ± 1 + | 144 ± 2 | 103 ± 2 | 101 ± 1 | 94 ± 1 |
| ∆*adhA* | 163 ± 1 | 141 ± 1 | ND | ND | ND |
| ∆*adhA*-∆*CC1311* | 165 ± 1 | 146 ± 2 | ND | ND | ND |
| $P_{zwf}$*C:T* | 192 ± 1 + | 150 ± 1 | 100 ± 1 | 98 ± 1 | 96 ± 1 |
| *adhA*$^{\Delta VQRY}$ $P_{zwf}$*C:T* | 134 ± 2 + | 149 ± 1 | 96 ± 1 | 95 ± 1 | 93 ± 2 |
| ∆*adhA* $P_{zwf}$*C:T* | 134 ± 1 | 149 ± 2 | ND | ND | ND |
| Evolved ~440 G [2 SNPs] | 157 ± 2 | ND | ND | ND | ND |
| *adhA*$^{\Delta VQRY}$ SNP2 | 162 ± 2 | ND | ND | ND | ND |
| Evolved ~1900 G [5 SNPs] | 118 ± 3 | ND | ND | ND | ND |
| *adhA*$^{\Delta VQRY}$ $P_{zwf}$*C:T* SNP3-4-5 | 126 ± 3 | ND | ND | ND | ND |
| **NA1000 background** | **M2G** | **M2X** | **PYE** | **PYEG** | **PYEX** |
| Parental | 129 ± 1 | ND | ND | ND | ND |
| $P_{zwf}$*T:C* | 166 ± 1 | ND | ND | ND | ND |

**Supplemental Table S1. Mutations in *adhA* (*CC1310*) and $P_{zwf}$ ($P_{CC2057}$) decreased the doubling time of the ancestor (CB15 ∆*pilA*).** Doubling time of *C. crescentus* CB15 ∆*pilA* and NA1000 derivative strains grown in minimal synthetic medium with glucose (M2G) or xylose (M2X) as a sole carbon source, or in PYE with or without glucose (PYEG) or xylose (PYEX). The two SNPs selected in the evolved population at ~440 generations (G) are *adhA*$^{\Delta VQRY}$ and *CC0212*$^{P77L}$. The five SNPs selected in the evolved population at ~1900 G are *adhA*$^{\Delta VQRY}$ (SNP1), *CC0212*$^{P77L}$ (SNP2) $P_{zwf}$*C:T* (SNP3) *CC1158*$^{W85*}$ (SNP4) and *rsaA*$^{W3G}$ (SNP5). The doubling time are expressed in min ± SD (n≥3).

| G1 proportion (%age) | | |
|---|---|---|
| **CB15 Δ*pilA* background** | **M2G** | **M2X** |
| Parental | 39.7 ± 0.6 | 30.1 ± 1 |
| *adhA*$^{\Delta VQRY}$ | 40.7 ± 0.3 | 30.8 ± 0.4 |
| Δ*adhA* | 41.6 ± 0.7 | ND |
| Δ*adhA*-Δ*CC1311* | 40.5 ± 0.7 | ND |
| $P_{zwf}$C:T | 28.9 ± 0.9 ** | 29.4 ± 0.9 |
| *adhA*$^{\Delta VQRY}$ $P_{zwf}$*C:T* | 30.5 ± 1.4 ** | 31.1 ± 0.6 |
| Δ*adhA* $P_{zwf}$*C:T* | 28.1 ± 0.5 ** | ND |
| **NA10000 background** | **M2G** | **M2X** |
| Parental | 33.2 ± 0.4 ** | 33.9 ± 0.5 |
| $P_{zwf}$*T:C* | 41.4 ± 0.5 | 34.0 ± 0.3 |

**Supplemental Table S2. Mutations in $P_{zwf}$ ($P_{CC2057}$) decrease the G1 lifetime of the ancestor (CB15 Δ*pilA*).** Percentage of G1 cells in *C. crescentus* CB15 Δ*pilA* and NA1000 derivative strains grown in minimal synthetic medium with glucose (M2G) or xylose (M2X) as a sole carbon source.

| Identifier | Strain/ Genotype | Source |
|---|---|---|
| SA001 | *NA1000-WT* | [1] |
| SA036 | *CB15 (ΔpilA)-adhA*$^{\Delta VQRY}$ *(CC1310)* | This study |
| SA037 | *CB15 (ΔpilA)-ΔadhA (CC1310)* | This study |
| SA038 | *CB15 (ΔpilA)*-$P_{zwf}$*C:T* | This study |
| SA039 | *CB15 (ΔpilA)- adhA*$^{\Delta VQRY}$ $P_{zwf}$*C:T* | This study |
| SA043 | *CB15 (ΔpilA)-WT* + plac290-$P_{zwf}$*C-lacZ* | This study |
| SA044 | *CB15 (ΔpilA)-*$P_{zwf}$*C:T* + plac290-$P_{zwf}$*T-lacZ* | This study |
| SA047 | *NA1000*-$P_{zwf}$*T:C* | This study |
| SA049 | *NA1000-WT* + plac290-$P_{zwf}$*T-lacZ* | This study |
| SA050 | *NA1000-*$P_{zwf}$*T:C* + plac290-$P_{zwf}$*C-lacZ* | This study |
| SA080 | *NA1000-WT* + pSKR478-$P_{lac}$-*rogfp2* | This study |
| SA081 | *NA1000-ΔfabH vanA::*$P_{vanA}$*-fabH* + pSKR478-$P_{lac}$-*rogfp2* | This study |
| SA082 | *NA1000-ΔpepC* | This study |
| UJ590 | *CB15-WT (ΔpilA)* | [2] |
| SA094 | *NA1000- idh::*$P_{vanA}$*-idh* | This study |
| SA097 | *NA1000*-$P_{zwf}$*T:C* + pSKR478-$P_{lac}$-*rogfp2* | This study |
| SA109 | *NA1000-ΔpepC* + pSKR478-$P_{lac}$-*pepC* | This study |
| IR037 | *NA1000-vanA::*$P_{vanA}$*-peredox_mCherry* | This study |
| RH944 | *CB15 (ΔpilA)-ΔadhA-ΔCC1311* | This study |
| RH756 | *CB15 (ΔpilA)- ΔadhA* $P_{zwf}$*C:T* | This study |
| RH46 | Evolved ~440 G [2 SNPs] | This study |
| RH179 | *adhA*$^{\Delta VQRY}$ SNP2 | This study |
| RH47 | Evolved ~1900 G [5 SNPs] | This study |
| RH684 | *adhA*$^{\Delta VQRY}$ $P_{zwf}$C:T SNP3-4-5 | This study |
| SM1131 | *NA1000-ΔfabH vanA::*$P_{vanA}$*-fabH* | [3] |

**Supplemental Table S3.** List of strains used in this study.

| Identifier | Description | Source |
|---|---|---|
| pSA025 | pSKR478-$P_{lac}$-*rogfp2* | This study |
| pSA035 | plac290-$P_{zwf}$*C-lacZ* | This study |
| pSA036 | plac290-$P_{zwf}$*C:T-lacZ* | This study |
| pSA037 | plac290-$P_{zwf}$*T-lacZ* | This study |
| pSA038 | plac290-$P_{zwf}$*T:C-lacZ* | This study |
| pSA027 | pSKR478-$P_{lac}$-*pepC* | This study |
| pIR001 | pVVENC2-*peredox-mCherry* | This study |
| pRH1 | pNPTS138 derivative carrying the in-frame deletion of *adhA*; $Kan^R$ | This study |
| pHG1 | pNPTS138 derivative carrying the in-frame deletion of *pepC*; $Kan^R$ | This study |
| pSKR410 | pVMCS2 derived carrying the in-frame deletion of *idh*; $Kan^R$ | This study |

**Supplemental Table S4.** List of plasmids used in this study.

| | Polar metabolites | | | | Non-Polar Metabolites | |
|---|---|---|---|---|---|---|
| | Non-derivatized polar metabolites | | Derivatized polar metabolites | | Lipids | |
| Resuspension solvent | 25% ACN + 5% ammonium acetate | | 1:1 (v/v) MeOH/$H_2O$ | | 2:1 (v/v) $CHCl_3$/MeOH | |
| Resuspension volume | 50 µL | | 50 µL | | 200 µL | |
| Loading volume | 10 µL | | 10 µL | | 10 µL | |
| Column | Fusion | | Fusion | | C18 | |
| Solvent A | 5 mM ammonium acetate in $H_2O$ | | 99.9% $H_2O$ + 0.1% FA | | 95:5 (v/v) $H_2O$/MeOH + 0.1% (v/v) ammonium hydroxide | |
| Solvent B | 100% ACN | | 99.9% MeOH + 0.1% FA | | 60:35:5 (v/v) IPA/MeOH/$H_2O$ + 0.1% (v/v) ammonium hydroxide | |
| Autosampler temperature | 10 °C | | 10 °C | | 10 °C | |
| Column oven temperature | 40 °C | | 40 °C | | 40 °C | |
| Flow rate | 0.5 mL/min | | | | 0.5 mL/min | |
| Gradient | %B | Time (min) | %B | Time (min) | %B | Time (min) |
| | 0 | 0 | 0 | 0 | 0 | 0 |
| | 5 | 3 | 5 | 3 | 0 | 4 |
| | 60 | 10 | 60 | 10 | 100 | 18 |
| | 95 | 11 | 95 | 11 | 100 | 25 |
| | 95 | 14 | 95 | 18 | 0 | 25.1 |
| | 5 | 15 | 60 | 19 | 0 | 30 |
| | 0 | 16 | 5 | 22 | | |
| | 0 | 21 | 0 | 25 | | |
| | | | 0 | 30 | | |

**Supplemental Table S5.** Details of the liquid-chromatography (LC) parameters used for performing LC-MS based experiments.

| | Polar metabolites | | Non-polar metabolites |
|---|---|---|---|
| | **Non-derivatized polar metabolites** | **Derivatized polar metabolites** | **Lipids** |
| Ionization mode | Negative | Positive | Negative |
| Drying Gas Temperature (°C) | 250 | 320 | 320 |
| Sheath Gas Temperature (°C) | 250 | 320 | 320 |
| Drying Gas Flow (L/min) | 10 | 10 | 10 |
| Sheath Gas Flow (L/min) | 10 | 10 | 10 |
| Nebulizer Pressure (psi) | 45 | 45 | 45 |
| Capillary Voltage (Vcap) | 4000 | 4000 | 4000 |
| Nozzle Voltage (V) | 1000 | 1000 | 1000 |
| Fragmentor Voltage (V) | 120 | 150 | 150 |
| Collision energy (eV) | 5, 15 | 5, 15 | 5, 15 |

**Supplemental Table S6.** Details of the Mass-spectrometry (MS) Parameters used for performing LC-MS based experiments.
